# An architectural RNA builds a reaction crucible through molecular concentration and activated enzyme selection

**DOI:** 10.64898/2026.11.08.765200

**Authors:** Tsuyoshi Ueno, Shungo Adachi, Tomohiro Yamazaki, Kensuke Ninomiya, Tetsuro Hirose

## Abstract

Architectural RNAs (arcRNAs) scaffold membraneless organelles (MLOs), yet how they organize functional proteins remains unclear. Nuclear stress bodies (nSBs), MLOs scaffolded by HSATIII arcRNAs, incorporate SRSFs during stress. During stress recovery, nSBs function as reaction crucibles that recruit the kinase CLK1 to accelerate SRSF rephosphorylation and normalize stress-induced splicing. Here, we identify the ASAP complex as a recruitment factor that recognizes Ser341-phosphorylated, catalytically activated CLK1. HSATIII-bound SRSFs recruit ASAP, assembling a platform for CLK1 recruitment. Biochemical reconstitution reveals that this platform enhances SRSF phosphorylation by concentrating SRSFs and CLK1 while selectively recruiting activated CLK1, establishing nSB reaction crucibles as molecular concentrators and selectors. The SRSF–ASAP–CLK1 interaction network is conserved between humans and mice, suggesting that primate-specific HSATIII arcRNAs co-opted this pre-existing network to establish reaction-crucible function in nSBs. Our findings reveal how arcRNAs organize functional proteins through quantitative enrichment and qualitative selection to build biochemically active MLOs.

## Introduction

Cells contain diverse membraneless organelles (MLOs) that accumulate specific proteins and nucleic acids without surrounding membranes.^1,2^ A subset of long noncoding RNAs (lncRNAs) serve as structural scaffolds that organize ribonucleoprotein condensates; we termed this class of scaffold lncRNAs architectural RNAs (arcRNAs).^3–6^ By recruiting and organizing proteins into distinct compartments, arcRNAs spatially and temporally coordinate diverse biological processes.^2,7^ Although many proteins associated with individual arcRNAs have been identified, the molecular principles by which arcRNAs selectively recruit functional proteins into MLO and organize them into biochemically active assemblies remain largely unknown.

Nuclear stress bodies (nSBs) are stress-induced MLOs assembled on HSATIII arcRNA.^8–12^ Upon thermal stress, HSF1 drives the transcription from primate-specific pericentromeric satellite III repeats, and the resulting HSATIII RNAs serve as arcRNAs that sequester multiple RNA-binding proteins to scaffold nSB assembly.^13–19^ HSATIII contains abundant GGAAUrepeats that concentrate a subset of SR splicing factors (SRSFs), including SRSF1, SRSF7, and SRSF9 within nSBs.^18,20–22^ Under normal conditions, SRSFs are highly phosphorylated, whereas thermal stress induces their global dephosphorylation and promotes stress-responsive splicing.^23–25^ Concomitantly, HSATIII sequesters dephosphorylated SRSFs into nSBs. During recovery, CLK1, a nuclear SRSF kinase, is selectively recruited to nSBs, where it rapidly rephosphorylates SRSFs and restores normal splicing (Extended Data Fig. 1a).^12,18^ Thus, nSBs undergo a functional transition from sequestration sites during thermal stress to reaction crucibles during recovery, where the local enrichment of SRSFs and CLK1 accelerates the SRSF rephosphorylation within 1 h after stress removal. Selective recruitment of CLK1 is therefore a key step that enables HSATIII to convert nSBs into biochemically active compartments.

We recently showed that recovery-specific recruitment of CLK1 depends on phosphorylation of Ser341, which is dynamically controlled by RIOK2 (RIO kinase 2) and PP1 (protein phosphatase 1).^26^ During thermal stress, PP1 dephosphorylates Ser341 and prevents CLK1 recruitment to nSBs, whereas during recovery, RIOK2 rephosphorylates Ser341, thereby enabling CLK1 recruitment. This phosphorylation cycle is coupled to temperature changes through the reversible thermosensor PPP1R2, which regulates PP1 activity in a temperature-dependent manner.^26^ These findings established Ser341 phosphorylation as a molecular switch for recovery-specific CLK1 recruitment but left unresolved how nSBs selectively recognize the phosphorylated CLK1 (Extended Data Fig. 1a).

Here, we identify the ASAP (apoptosis- and splicing-associated protein) complex^27^ as a phospho-selective recruiting factor that recognizes Ser341-phosphorylated, enzymatically activated CLK1 and recruits it to nSBs. ASAP multivalently links CLK1 to SRSFs concentrated on the GGAAU repeats of HSATIII arcRNA through multiple binding sites for both CLK1 and SRSFs. Biochemical reconstitution demonstrates that these interactions enhance SRSF phosphorylation through quantitative enhancement of enzyme-substrate association and qualitative selection of Ser341-phosphorylated, enzymatically activated CLK1. Our findings reveal how arcRNAs organize functional reaction crucibles.

## Results

### ASAP is a promising candidate recruiting factor for phosphorylated CLK1

We previously showed that CLK1 Ser341 phosphorylation by RIOK2 and the nSB- localization regions (nSB-LRs) within the CLK1 intrinsically disordered region (IDR) are both required for its localization to nSBs (Extended Data Fig. 1b).^26^ We hypothesized that phosphorylation-dependent CLK1 localization to nSBs is mediated byrecruitment factors that specifically recognize Ser341-phosphorylated CLK1. We sought proteins that (i) interact with CLK1 in a Ser341 phosphorylation-dependent manner, (ii) bind to CLK1 via the nSB-LRs,^26^ (iii) reside in nSBs, and (iv) are required for CLK1 localization to nSBs. For the first two criteria, we performed co-immunoprecipitation (co-IP) coupled with mass spectrometry (MS) using two complementary screens to identify proteins whose interaction with CLK1 was abolished either by inhibition of Ser341 phosphorylation with the RIOK2 inhibitor CQ211^28^ or by deletion of the nSB-LRs (Extended Data Fig. 1b,c).^26^ The intersection of these screens identified 26 candidates, including components of two related complexes, ASAP (ACIN1-RNPS1-SAP18) and PSAP (PNN-RNPS1-SAP18) (Fig. 1a, Extended Data Fig. 1d-f and Supplementary Table 1). Among the candidates, ACIN1 and RNPS1 were showing marked reductions in CLK1 interaction under both conditions relative to their respective control. Co-IP followed by western blotting (WB) validated these interactions and their dependence on RIOK2 activity and nSB-LRs (Fig. 1b,c). To eliminate effects of CLK1 kinase activity, all subsequent binding and localization analyses in this study were performed using the kinase-inactive mutant CLK1 K191R. We confirmed that CLK1 K191R exhibited nSB localization dynamics comparable to those of wild-type CLK1.^12,26^

**Figure 1.**
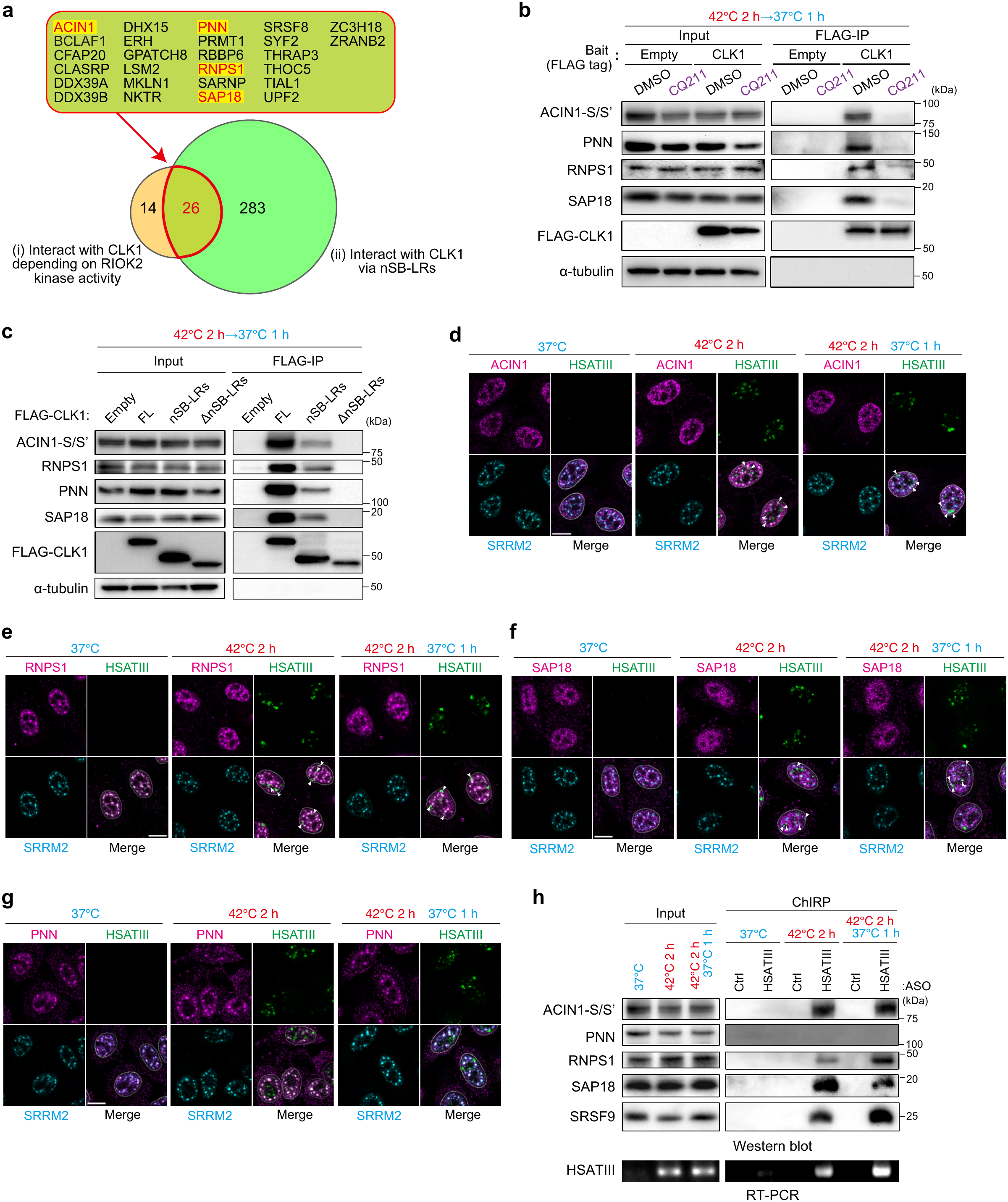
The ASAP complex is a candidate recruiting factor for phosphorylated CLK1. (a) Venn diagram comparing proteins preferentially interacting with CLK1 in the absence versus presence of the RIOK2 inhibitor CQ211 (Interacting with CLK1 depending on RIOK2 kinase activity) or with full-length CLK1 versus the nSB-LR deletion mutant (Interacting with CLK1 via nSB-LRs), identified by coIP-MS (log_2_ fold change < -1.0, *P* < 0.05 (two-tailed t test)). The 26 overlapping proteins are listed above. Co-IP assays were performed using whole-cell lysates from HeLa cells subjected to thermal stress recovery (42°C 2 h → 37 °C 1 h). CQ211 (10 μM) was added during recovery. (b) Co-IP of FLAG-CLK1 with the indicated ASAP and PSAP components during recovery in the presence or absence of the RIOK2 inhibitor CQ211. Co-IP was performed using whole-cell lysates from HeLa cells subjected to thermal stress recovery (42°C 2 h → 37 °C 1 h). CQ211 (10 μM) was added during recovery. α -tubulin was used as a negative control for CLK1 co-IP. (c) Co-IP of CLK1 nSB-LR deletion mutants with the indicated ASAP and PSAP components. Co-IP was performed using whole-cell lysates from HeLa cells subjected to thermal stress recovery (42°C 2 h → 37 °C 1 h). α-tubulin was used as a negative control for CLK1 co-IP. (d-g) Localization of endogenous ACIN1 (d), RNPS1 (e), SAP18 (f), and PNN (g) to HSATIII-scaffolded nSBs and SRRM2-marked nuclear speckles under the indicated temperature conditions. Nuclei were stained with DAPI. Scale bars, 10 μm. Arrowheads indicate colocalization with nSBs. (h) HSATIII ChIRP followed by Western blot for the indicated ASAP and PSAP components using HSATIII-targeting or random control antisense oligonucleotides (ASOs) under the indicated temperature conditions. SRSF9 was used as a positive control for HSATIII-ChIRP.

ASAP and PSAP share RNPS1 and SAP18 but contain ACIN1 and PNN as their complex-specific subunits, respectively (Extended Data Fig. 1d).^29^ These exon junction complex (EJC)-associated complexes function in multiple aspects of RNA metabolism, including pre-mRNA splicing, mRNA export, and nonsense-mediated mRNA decay,^29–34^ but no functional link between either complex and CLK1 has been reported to date.

We next examined whether ASAP and PSAP satisfy the third criterion by combining HSATIII RNA fluorescence in situ hybridization (RNA-FISH) with immunofluorescence (IF). ACIN1, RNPS1, and SAP18 were broadly distributed throughout the nucleoplasm and colocalized with nSBs during and after thermal stress, whereas PNN showed no detectable nSB localization (Fig. 1d-g). All four proteins localized to nuclear speckles under all temperature conditions, consistent with previous reports.^27,30,35^ HA-tagged proteins showed more prominent enrichment in nuclear speckles but retained nSB localization patterns consistent with the endogenous proteins, with PNN again showing no detectable nSB localization (Extended Data Fig. 1g-k). Because ACIN1 undergoes extensive alternative splicing, generating multiple protein isoforms including L, S, and S′, we analyzed the ACIN1-S isoform, which has been reported to assemble into the ASAP complex (Extended Data Fig. 1k).^27,36^ Functional analyses of other ACIN1 isoforms are described below. To further evaluate ASAP and PSAP localization to nSBs, we performed ChIRP (chromatin isolation by RNA purification) targeting HSATIII arcRNA, with SRSF9 serving as a positive control. ACIN1, RNPS1, and SAP18, but not PNN, were recovered during and after thermal stress (Fig. 1h), independently confirming the selective association of ASAP with nSBs. Thus, ASAP, but not PSAP, satisfies the third criterion for a phosphorylated CLK1-recruitment factor.

### ASAP facilitates nSB-mediated splicing regulation by promoting CLK1 localization

To examine the fourth criterion, we analyzed CLK1 localization to nSBs following knockdown of ASAP components. Knockdown of ACIN1 using either an ACIN1-L-specific siRNA (#1) or an siRNA targeting all ACIN1 isoforms (L, S, and S′; #2) significantly impaired CLK1 localization to nSBs (Fig. 2a,b and Extended Data Fig. 2a). RNPS1 knockdown similarly reduced CLK1 localization (Fig. 2c,d and Extended Data Fig. 2b). Knockdown of either ACIN1 or RNPS1 did not substantially affect expression of the other protein (Extended Data Fig. 2c). In contrast, knockdown of the PSAP-specific component PNN had no significant effect (Extended Data Fig. 2d-f). We next examined whether two additional candidates, THRAP3 and BCLAF1, which satisfied the first three screening criteria (Fig. 1a and our previous study^12^), also contributed to CLK1 localization or not. Double knockdown of these functionally redundant proteins^37^ did not affect CLK1 localization (Extended Data Fig. 2g-i), further supporting a specific role for ASAP. CLK1 localizes to nuclear speckles independently of Ser341 phosphorylation,^12,26^ and depletion of ACIN1 or RNPS1 did not affect this localization (Extended Data Fig. 2j,k), indicating that ASAP specifically promotes CLK1 recruitment to nSBs rather than its general localization to nuclear compartments. Re-expression of either ACIN1-L or ACIN1-S restored CLK1 localization in cells depleted of all ACIN1 isoforms (Extended Data Fig. 2l,m), and exogenous RNPS1 similarly rescued the effect of RNPS1 knockdown (Extended Data Fig. 2n,o), confirming the specificity of these knockdown phenotypes.

**Figure 2.**
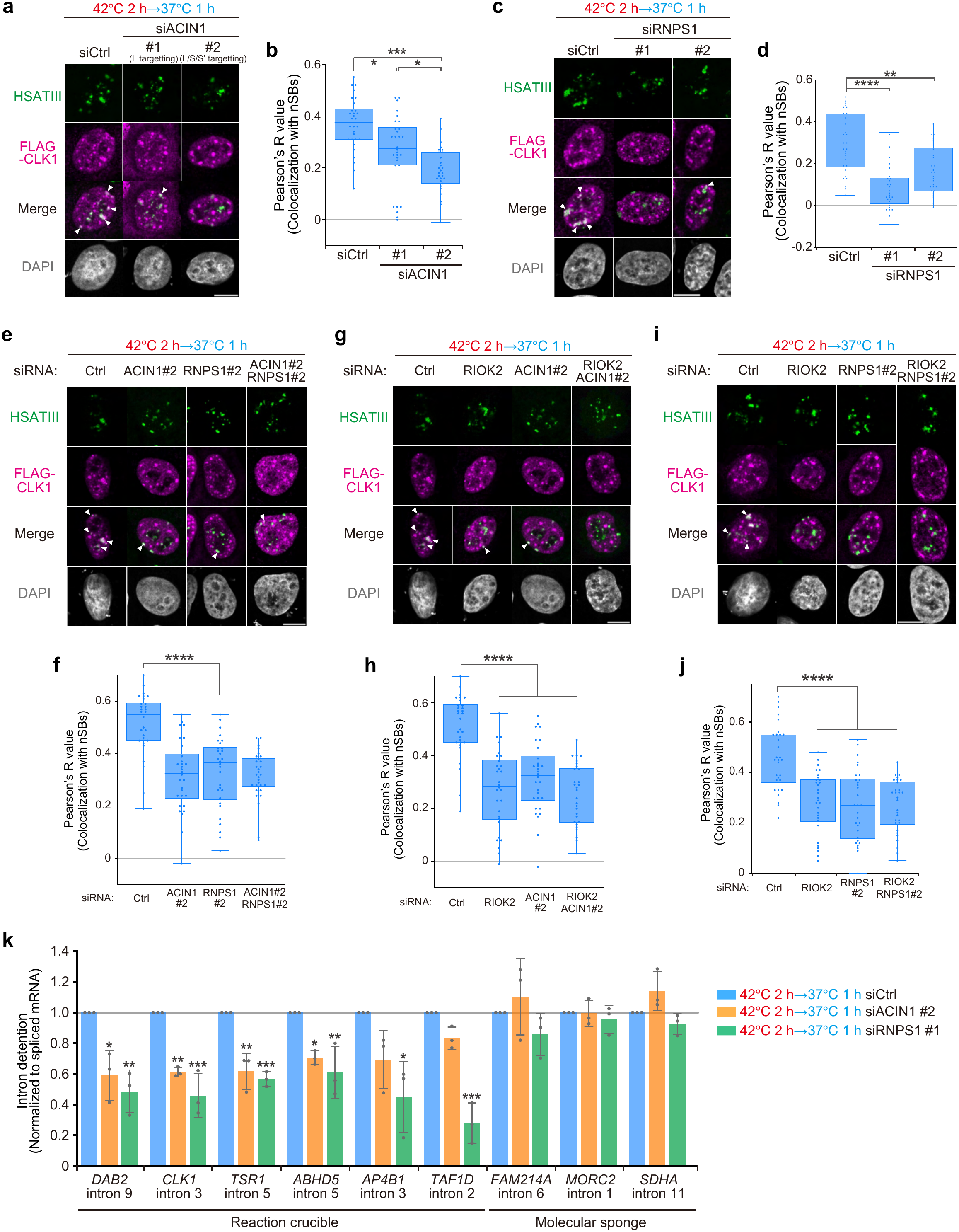
The ASAP complex is required for CLK1 localization to nSBs and nSB- mediated splicing regulation. (a, c) CLK1 localization to nSBs during stress recovery following knockdown of ACIN1 (a) or RNPS1 (c). siACIN1 #1 targets the ACIN1-L isoform, and siACIN1 #2 targets all ACIN1 isoforms (a). Nuclei were stained with DAPI. Scale bars, 10 μm. siRNA, 20 nM. Arrowheads indicate CLK1 colocalization with nSBs. (b, d, f, h, j) Quantification of colocalization in (a), (c), (e), (g), and (i), respectively using Pearson’s correlation coefficients (R) between CLK1 and HSATIII within nuclei. n = 30. Kruskal-Wallis test followed by Dunn’s multiple-comparisons test was used. **** *P* < 0.0001, *** *P* < 0.001, ** *P* < 0.01, * *P* < 0.05. (e, g, i) CLK1 localization to nSBs during stress recovery following combined knockdown of ACIN1 and RNPS1 (e), RIOK2 and ACIN1 (g), or RIOK2 and RNPS1 (i). For single knockdown, siRNA was used at 20 nM; for double knockdown, each siRNA was used at 10 nM. (k) RT-qPCR analysis of intron detention in the pre-mRNA of *DAB2, CLK1, TSR1, ABHD5, AP4B1,* and *TAF1D* (Reaction crucible-dependent targets), and *FAM214A, MORC2,* and *SDHA* (Molecular sponge-dependent targets) following ACIN1 or RNPS1 knockdown during thermal stress recovery. Intron detention levels were normalized to the corresponding spliced RNAs to account for changes in target gene expression. Data are shown as mean ± SD. n = 3. Šídák’s multiple-comparisons test: **** *P* < 0.0001, *** *P* < 0.001, ** *P* < 0.01, * *P* < 0.05.

To test whether ACIN1 and RNPS1 function in the same pathway to regulate CLK1 localization, we performed double knockdown of both proteins. Double knockdown of ACIN1 and RNPS1 impaired CLK1 localization to a similar extent as either single knockdown (Fig. 2e,f). To determine whether RIOK2-mediated phosphorylation and ASAP-dependent recruitment act in the same pathway, we combined RIOK2 depletion with depletion of either ACIN1 or RNPS1. Combined depletion of RIOK2 with either ACIN1 or RNPS1 produced no additional reduction in CLK1 localization (Fig. 2g-j). These results support a model in which RIOK2-mediated phosphorylation and ASAP-dependent recruitment act sequentially in the same pathway to recruit CLK1 to nSBs.

We next examined whether impaired CLK1 recruitment affects the reaction-crucible activity of nSBs. Because nSB-localized CLK1 accelerates SRSF rephosphorylation,^12^ we first examined SRSF phosphorylation during the early phase of stress recovery. ACIN1 and RNPS1 depletion reduced the phosphorylation of several SRSFs detected by the pan-phospho-SRSF antibody 1H4 during early recovery, whereas no clear difference was observed during late recovery (Extended Data Fig. 2p,q). These results indicate that ASAP accelerates a specific subset of SRSF rephosphorylation during stress recovery, as knockdown of ACIN1 and RNPS1 delayed this process. We previously showed that nSBs regulate intron detention through two distinct mechanisms^12,18,22^: a molecular sponge mechanism that sequesters m^6^A modification-related factors and suppresses m^6^A-dependent splicing, and a reaction crucible mechanism that promotes CLK1-dependent SRSF phosphorylation and SRSF-dependent intron detention. Similar to RIOK2 inhibition,^26^ knockdown of ACIN1 or RNPS1 selectively impaired reaction crucible-dependent splicing without affecting molecular sponge-dependent splicing (Fig. 2k). Together, these findings demonstrate that ASAP promotes CLK1 recruitment to nSBs and thereby accelerates SRSF rephosphorylation and reaction crucible-dependent splicing during stress recovery, thus satisfying all four criteria for a phosphorylated CLK1-recruitment factor.

### ASAP directly and selectively binds to Ser341-phosphorylated CLK1

Because ACIN1 and RNPS1 depletion caused similar defects in CLK1 localization, we asked whether assembly of an intact ASAP complex is required for efficient CLK1 binding. Depletion of either ACIN1 or RNPS1 significantly reduced the interaction between CLK1 and the remaining ASAP components (Fig. 3a,b). Consistently, disruption of ASAP assembly by deletion of the ACIN1 RSB (RNPS1/SAP18-binding) domain or the RNPS1 RRM (RNA recognition motif) domain^29^ impaired CLK1 binding (Fig. 3c-h), and neither mutant rescued CLK1 localization following knockdown (Extended Data Fig. 3a-f). These results indicate that intact ASAP complex assembly is required for efficient CLK1 binding and recruitment to nSBs.

**Figure 3.**
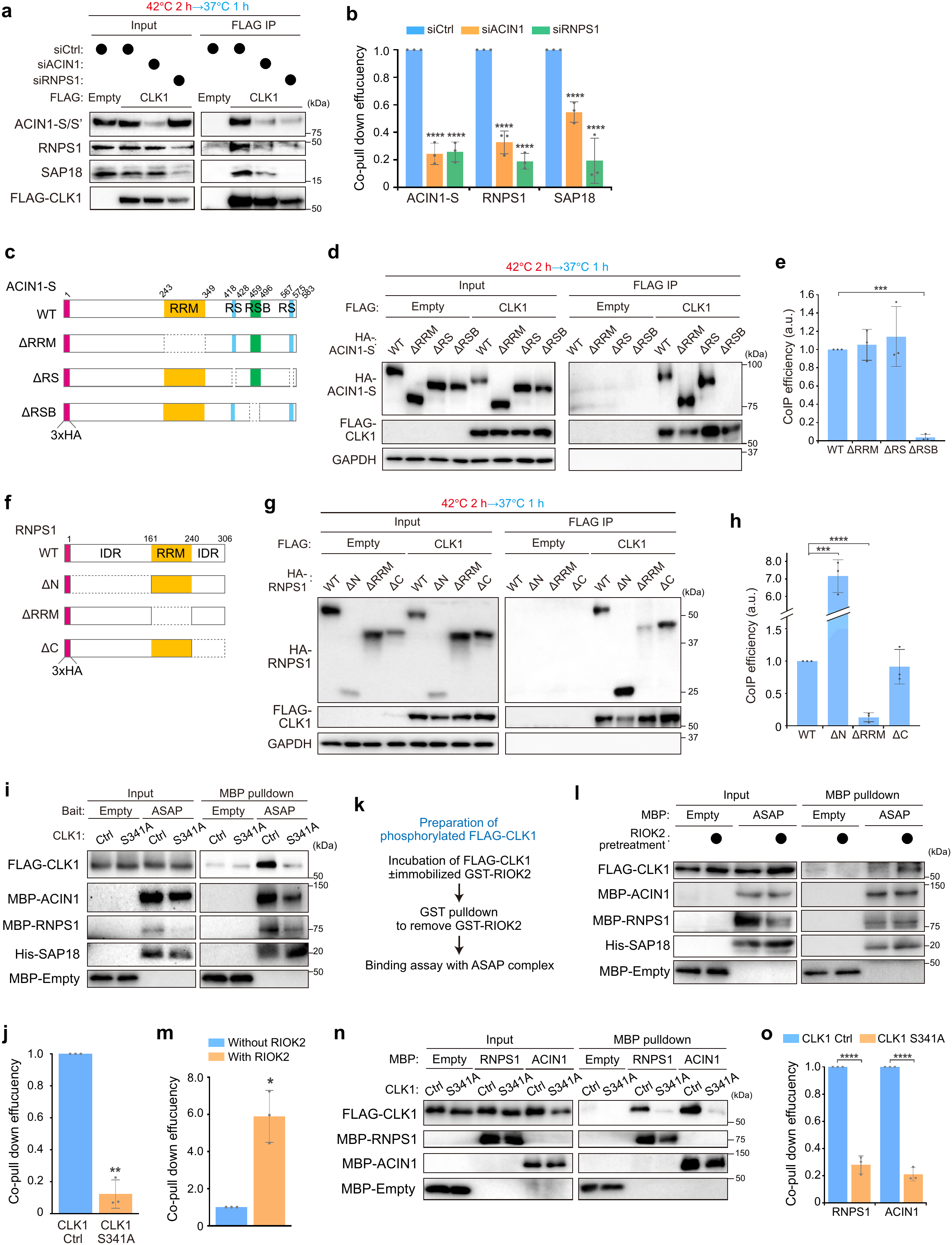
The ASAP complex directly and selectively binds to Ser341-phosphorylated CLK1. (a, b) Co-IP of FLAG-CLK1 with ASAP components following ACIN1 or RNPS1 knockdown (a) and quantification of co-IP efficiency (b), calculated as the ratio of ASAP components to CLK1. Šídák’s multiple-comparisons test. **** *P* < 0.0001. siRNA, 20 nM. Data are shown as mean ± SD. n = 3. Co-IP was performed using whole-cell lysates from HeLa cells subjected to thermal stress recovery (42°C 2 h → 37 °C 1 h). (c–h) Interaction mapping of CLK1 with ACIN1 and RNPS1. (c, f) Schematic representation of ACIN1 and RNPS1 deletion mutants. RRM, RNA recognition motif; RS, serine-arginine-rich region; RSB, RNPS1-SAP18 binding domain; IDR, intrinsically disordered region. (d, g) Co-IP of FLAG-CLK1 with ACIN1 (d) or RNPS1 (g) deletion mutants. (e, h) Quantification of co-IP efficiency in (d) and (g), calculated as the ratio of ACIN1 or RNPS1 to CLK1. GAPDH was used as a negative control for CLK1 co-IP. Šídák’s multiple-comparisons test. **** *P* < 0.0001, *** *P* < 0.001. Data are shown as mean ± SD. n = 3. Co-IP was performed using whole-cell lysates from HeLa cells subjected to thermal stress recovery (42°C 2 h → 37 °C 1 h). (i, j) In vitro binding of recombinant ASAP to Control (Ctrl) or S341A CLK1 (i) and quantification of co-pulldown efficiency (j), calculated as the ratio of CLK1 to ASAP components. Two-tailed t test was used. ** *P* < 0.01. Data are shown as mean ± SD. n = 3. (k-m) In vitro binding of ASAP to CLK1 preincubated with or without RIOK2. (k) Workflow for the preparation of phosphorylated CLK1. (l) In vitro binding of recombinant ASAP to CLK1 with or without RIOK2-mediated phosphorylation. (m) Quantification of co-pulldown efficiency in (l), calculated as the ratio of CLK1 to ASAP components. Two-tailed t test was used. * *P* < 0.05. Data are shown as mean ± SD. n = 3. (n, o) In vitro binding of RNPS1 and ACIN1 to control and S341A CLK1 recombinant proteins (n) and quantification of co-pulldown efficiency (o), calculated as the ratio of CLK1 to RNPS1 or ACIN1. Data are shown as mean ± SD. n = 3. Šídák’s multiple-comparisons test; **** *P* < 0.0001.

We next asked whether ASAP directly and selectively recognizes Ser341-phosphorylated CLK1. To eliminate effects of CLK1 kinase activity, we prepared recombinant CLK1 proteins carrying the kinase-dead CLK1 K191R mutation as the control (Ctrl), and an additional S341A mutation in the same background. Because both proteins were purified from HEK293 cells (Extended Data Fig. 3g), their key difference was Ser341 phosphorylation status. To test whether ASAP directly recognizes Ser341-phosphorylated CLK1, we reconstituted the ASAP complex from recombinant MBP-RNPS1, ACIN1 and His-SAP18 (Extended Data Fig. 3g). The resulting complex was purified by sequential His- and MBP-affinity purification. The reconstituted ASAP complex directly bound Ctrl CLK1 but showed substantially reduced binding to the S341A mutant (Fig. 3i,j). In contrast, SRSF9, a CLK1 substrate, showed no binding preference between Ctrl and S341A CLK1 (Extended Data Fig. 3h,i). To directly test whether Ser341 phosphorylation enhances ASAP binding, we phosphorylated recombinant CLK1 purified from E. coli in vitro using RIOK2 and examined its binding to ASAP (Fig. 3k and Extended Data Fig. 3g,j). RIOK2-mediated phosphorylation markedly enhanced CLK1 binding to ASAP (Fig. 3l,m), establishing a direct link between CLK1 Ser341 phosphorylation and ASAP recognition.

We then examined which ASAP subunits mediate this interaction. Recombinant CLK1 directly bound to ACIN1 and RNPS1, but not SAP18 (Extended Data Fig. 3k,l), and these interactions were inhibited by the S341A mutation (Fig. 3n,o). Thus, ACIN1 and RNPS1 provide two separable Ser341-dependent binding surfaces for CLK1, together supporting efficient CLK1 binding to the intact ASAP complex in cells. Together, these findings demonstrate that ASAP directly and selectively recognizes Ser341-phosphorylated CLK1 through multiple interactions involving ACIN1 and RNPS1, establishing a molecular basis for phosphorylation-dependent CLK1 recruitment to nSBs.

### Artificial tethering of ASAP recruits Ser341-phosphorylated CLK1 in cells

To determine whether tethering ASAP to an ectopic RNA scaffold is sufficient to recruit CLK1 in cells, we established an ectopic RNA scaffold system. In this system, MCP (MS2 coat protein)-fused ACIN1, a specific component of ASAP, was tethered to an engineered RNA containing tandem MS2 stem loops (Fig. 4a). MCP-fused ACIN1 tethered to MS2 RNA formed discrete nuclear RNA foci that colocalized with CLK1 (Fig. 4b,c). These ectopic foci did not colocalize with nuclear speckles (Extended Data Fig. 4a,b), indicating that CLK1 recruitment to ACIN1-tethered RNA foci does not result from their association with nuclear speckles. Because nSBs are absent under the normal temperature conditions used here, these results demonstrate that artificial tethering of ACIN1 can recruit CLK1 independently of endogenous nSBs or nuclear speckles.

**Figure 4.**
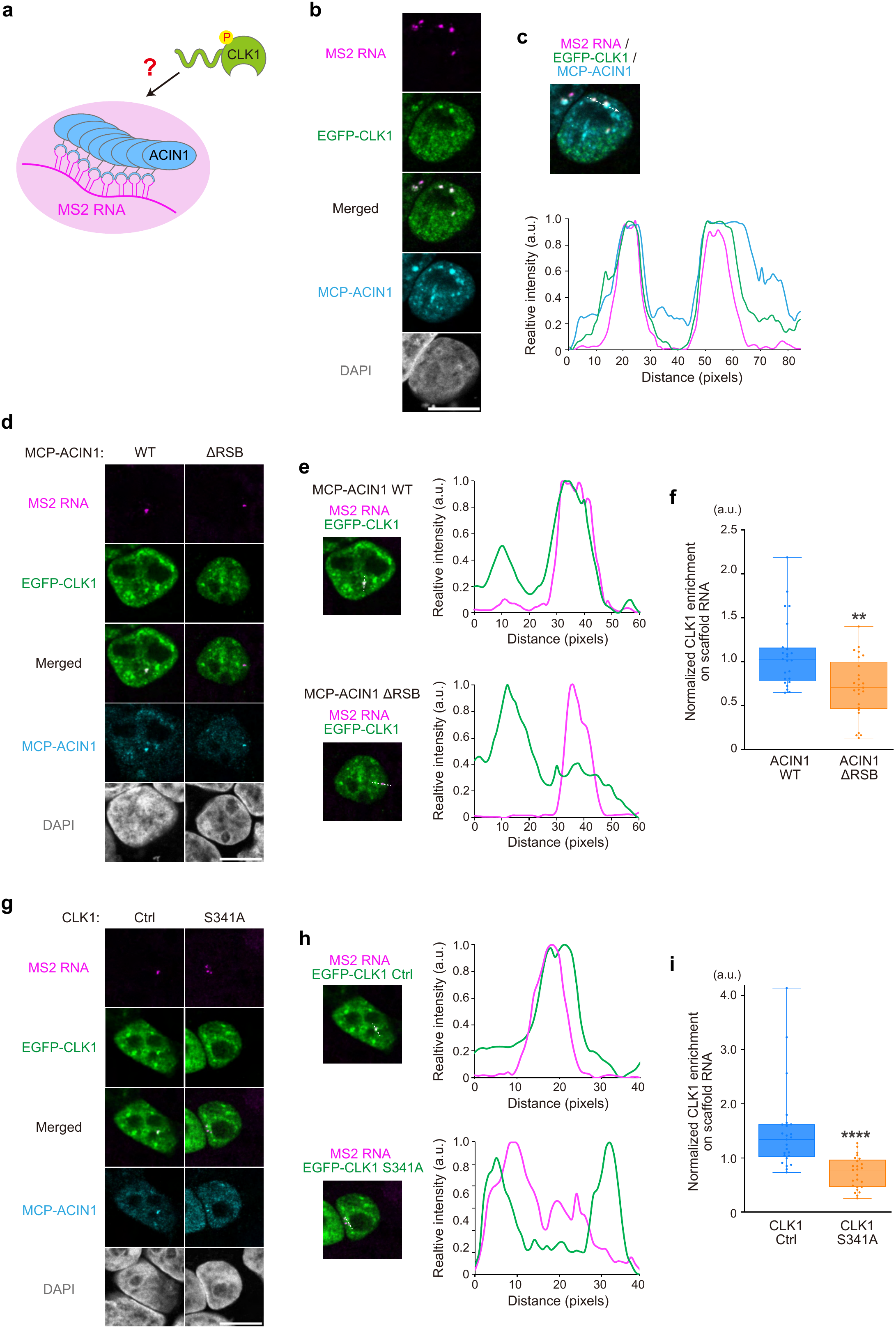
ASAP recruits Ser341-phosphorylated CLK1 in cells. (a) Schematic of the MS2 tethering assay used to evaluate ASAP-mediated CLK1 recruitment. The scaffold RNA was visualized using PP7 stem-loop sequences. (b, c) Localization of EGFP-CLK1 to MCP-ACIN1-tethered MS2 RNA foci (b) and line-scan profiles of MS2 RNA (magenta), EGFP-CLK1 (green), MCP-ACIN1 (blue) fluorescence along the indicated line (c). Nuclei were stained with DAPI. Scale bars, 10 μm. (d, g) Localization of EGFP-CLK1 to MS2 RNA tethered with MCP-ACIN1 WT or ΔRSB (d), or of EGFP-CLK1 Control or S341A to MCP-ACIN1-tethered MS2 RNA (g). (e, h) Line-scan profiles of MS2 RNA (magenta) and EGFP-CLK1 (green) fluorescence along the indicated lines in (d) and (g). (f, i) Quantification of CLK1 recruitment in (d) and (g). CLK1 intensity within MS2 RNA foci was normalized to nuclear CLK1 intensity and MS2 RNA focus intensity. n = 25. Mann-Whitney U test was used. **** *P* < 0.0001, ** *P* < 0.01.

We next examined whether efficient CLK1 recruitment requires ASAP complex assembly. RNPS1 and SAP18 also colocalized with ACIN1 foci, consistent with assembly of the ASAP complex on the ectopic RNA scaffold (Extended Data Fig. 4c). The ACIN1 ΔRSB mutant, which is defective in ASAP assembly,^29^ showed markedly reduced CLK1 recruitment to the foci (Fig. 4d-f). Likewise, RNPS1 knockdown impaired CLK1 recruitment to ACIN1 foci (Extended Data Fig. 4d,e). These results indicate that assembly of the ASAP complex is required for efficient CLK1 recruitment in cells.

Finally, we examined whether CLK1 recruitment to ACIN1 foci depends on Ser341 phosphorylation. Recruitment of the CLK1 S341A mutant to ACIN1 foci was markedly reduced compared with CLK1 Ctrl (Fig. 4g-i), and RIOK2 inhibition, which reduces Ser341 phosphorylation, similarly impaired CLK1 recruitment (Extended Data Fig. 4f,g). Thus, ASAP recognition of Ser341-phosphorylated CLK1 is also evident in cells. Together, these results demonstrate that the assembled ASAP complex recruits Ser341-phosphorylated CLK1 in cells.

### The HSATIII-SRSF-ASAP complex forms an RNA-based recruitment platform for Ser341-phosphorylated CLK1 in nSBs

ASAP recruits CLK1 to HSATIII-scaffolded nSBs. To define the molecular interactions underlying this recruitment, we established an in vitro RNA pulldown assay to reconstitute the recruitment of Ser341-phosphorylated CLK1 to HSATIII-scaffolded complex. In this system, biotinylated GGAAU-repeats, mimicking HSATIII, were incubated with purified recombinant proteins and recovered using streptavidin beads (Extended Data Fig. 5a). ASAP alone exhibited little affinity for GGAAU-repeats, indicating that additional factors mediate its recruitment to HSATIII arcRNA (Fig. 5a). Because SRSFs are highly enriched in nSBs and serve as substrates in the reaction crucible function, we hypothesized that SRSFs serve as adaptors to recruit ASAP onto HSATIII arcRNA. Indeed, SRSF9 directly bound GGAAU-repeats and efficiently recruited ASAP to GGAAU-repeats (Fig. 5b). Similar results were obtained with SRSF1 (Extended Data Fig. 5b,c); therefore, we used SRSF9, one of the SRSFs most prominently enriched in nSBs,^12^ as a representative SRSF for subsequent analyses. SRSF9 directly bound to all three ASAP components in vitro (Extended Data Fig. 5d), suggesting multivalent interaction between SRSF9 and the ASAP complex. ASAP-SRSF9 interaction was confirmed in cells by conducting ACIN1 co-IP (Extended Data Fig. 5e,f). To determine whether SRSF9 also promotes ASAP recruitment to endogenous nSBs, we depleted SRSF9 and examined ASAP association with HSATIII by conducting HSATIII-ChRIP. SRSF9 depletion markedly reduced ASAP association with endogenous HSATIII, whereas recruitment of SAFB, another nSB marker protein, was unaffected (Fig. 5c,d). Thus, SRSFs act as adaptors that recruit ASAP to HSATIII arcRNA scaffold.

**Figure 5.**
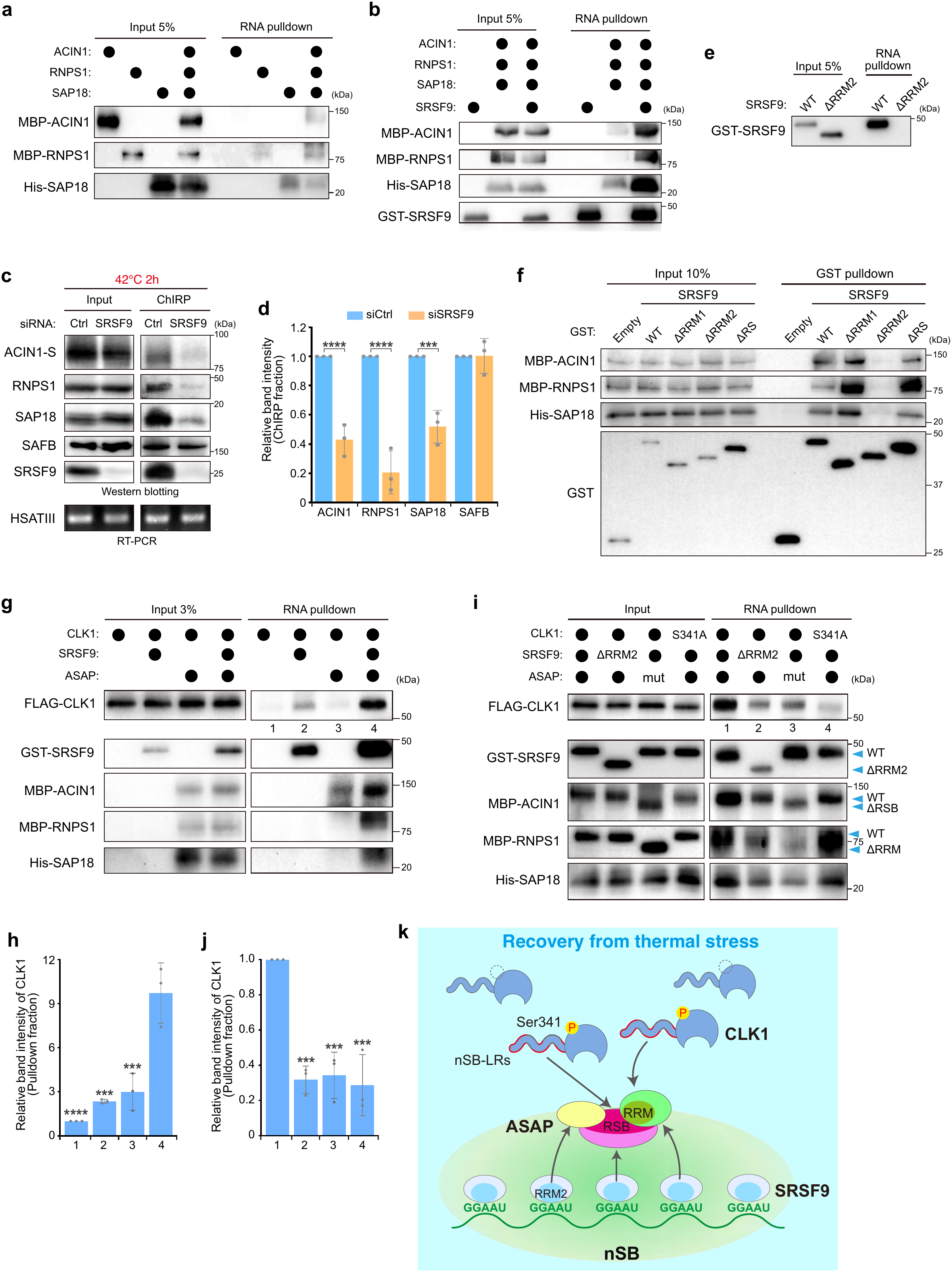
The HSATIII-SRSF-ASAP assembly recruits phosphorylated CLK1 in nSBs. (a, b, e) In vitro RNA pulldown assays using the indicated recombinant proteins: (a) individual ASAP components or the ASAP complex; (b) SRSF9, ASAP, or their combination; and (e) SRSF9 WT and ΔRRM2. The proteins were incubated with biotinylated GGAAU-repeat RNA, followed by pulldown using streptavidin beads. (c, d) HSATIII ChIRP of ASAP association with nSBs following SRSF9 knockdown (c) and quantification of band intensities in the ChIRP fraction (d). Šídák’s multiple-comparisons test was used. **** *P* < 0.0001, *** *P* < 0.001. Data are shown as mean ± SD. n = 3. (f) In vitro binding of SRSF9 WT and deletion mutants to the ASAP complex. Recombinant GST-tagged SRSF9 WT and deletion mutants were incubated with the in vitro-assembled ASAP complex and subjected to GST pulldown. (g, i) In vitro RNA pulldown assay to evaluate CLK1 recruitment to GGAAU-repeat RNA in the presence of the indicated components (g), and following disruption of the interaction surfaces within the HSATIII-SRSF-ASAP-CLK1 platform (i). RNA pulldown was performed as in (a). (h, j) Quantification of CLK1 band intensity in (g) and (i). Šídák’s multiple-comparisons test was used. **** *P* < 0.0001, *** *P* < 0.001. Data are shown as mean ± SD. n = 3. (k) Schematic model of the HSATIII-SRSF-ASAP platform mediating CLK1 recruitment to nSBs. During thermal stress, SRSF9 is sequestered on the repetitive GGAAU sequences of HSATIII arcRNA through multivalent interactions. SRSF9 binds HSATIII RNA through its RRM2 domain, which also mediates interactions with all three ASAP components, thereby assembling an HSATIII-SRSF-ASAP platform. During recovery, the ASAP complex selectively recognizes Ser341-phosphorylated CLK1 through multivalent interactions of ACIN1 and RNPS1 with nSB-localization regions (nSB-LRs) within the CLK1 IDR, thereby promoting CLK1 recruitment to HSATIII-scaffolded nSBs.

To define the molecular basis of the adaptor function, we next mapped the SRSF9 domain responsible for this function. Because RRM2 of SRSF1 is required for HSATIII binding and nSB localization (Extended Data Fig. 5g),^38^ we examined the corresponding domain of SRSF9. Deletion of RRM2 abolished both HSATIII binding and SRSF9 localization to nSBs (Fig. 5e and Extended Data Fig. 5h,i). Consistently, the isolated RRM2 domain bound GGAAU-repeats substantially more efficiently than RRM1 (Extended Data Fig. 5j-l). Additionally, RRM2 deletion also abolished binding to all three ASAP components (Fig. 5f and Extended Data Fig. 5m). Thus, RRM2 is required for both HSATIII binding and interactions with all three ASAP components, providing the molecular basis for SRSF9-mediated recruitment of ASAP to HSATIII.

We then tested whether the HSATIII-SRSF-ASAP complex function as a platform for phosphorylated CLK1 recruitment. CLK1 alone exhibited little binding to GGAAU-repeats, and addition of either SRSF9 or ASAP alone resulted in limited recruitment (Figure 5g,h). In contrast, combining SRSF9 and ASAP substantially enhanced recruitment of phosphorylated CLK1 to GGAAU-repeats (Fig. 5g,h), recapitulating the mechanism of phosphorylated CLK1 recruitment to nSBs in vitro. To test whether this recruitment requires each interaction surface within the platform, we used SRSF9 ΔRRM2, an ASAP mutant complex containing ACIN1 ΔRSB and RNPS1 ΔRRM (ASAP mut), and CLK1 S341A, perturbing SRSF-mediated assembly on HSATIII, ASAP complex integrity, and phosphorylation-dependent CLK1 recognition, respectively (Extended Data Fig. 5n). Each perturbation markedly impaired CLK1 recruitment to GGAAU-repeats (Fig. 5i,j), demonstrating that efficient CLK1 recruitment by the HSATIII-SRSF-ASAP platform depends on an intact multivalent interaction network. Together, these results establish the HSATIII-SRSF-ASAP complex as a molecular platform for the selective recruitment of Ser341-phosphorylated CLK1. Efficient recruitment requires intact ASAP complex assembly and is mediated by multivalent protein-protein interactions of ASAP with both phosphorylated CLK1 and SRSF9, together with multivalent RNA-protein interactions between SRSF9 and the repetitive GGAAU sequences of HSATIII arcRNA (Fig. 5k).

### The HSATIII-SRSF-ASAP platform enhances SRSF phosphorylation through quantitative enrichment and qualitative selection of activated CLK1

We next examined the functional significance of the HSATIII-SRSF-ASAP platform in the crucible function of nSBs. Although SRSFs can already bind CLK1 directly as substrates, we asked what additional functional advantage this platform provides. We first examined whether the platform quantitatively enhances CLK1-SRSF9 association. Consistent with efficient recruitment of CLK1 to the platform (Fig. 5g), CLK1 association with SRSF9 was highest when all components were present. Omitting either GGAAU or ASAP reduced this association to approximately 20-30% of that level (Fig. 6a,b). We next asked whether this enhanced association promotes SRSF phosphorylation. We reconstituted SRSF9 phosphorylation in vitro using GGAAU-repeat RNA and recombinant SRSF9, ASAP, and catalytically active CLK1 WT.^26^ Following incubation with ^32^P-labeled ATP, SRSF9 was isolated by GST pulldown and its phosphorylation level was assessed by autoradiography (Extended Data Fig. 6a). The HSATIII-SRSF-ASAP platform enhanced SRSF9 phosphorylation, whereas omission of either GGAAU or ASAP reduced phosphorylation to 50% or 30%, respectively (Fig. 6c,d). The HSATIII-SRSF-ASAP platform enhanced SRSF9 phosphorylation by more than 4-fold compared with SRSF9 and CLK1 alone, indicating that the platform quantitatively promotes enzyme-substrate association and CLK1-mediated SRSF phosphorylation.

**Figure 6.**
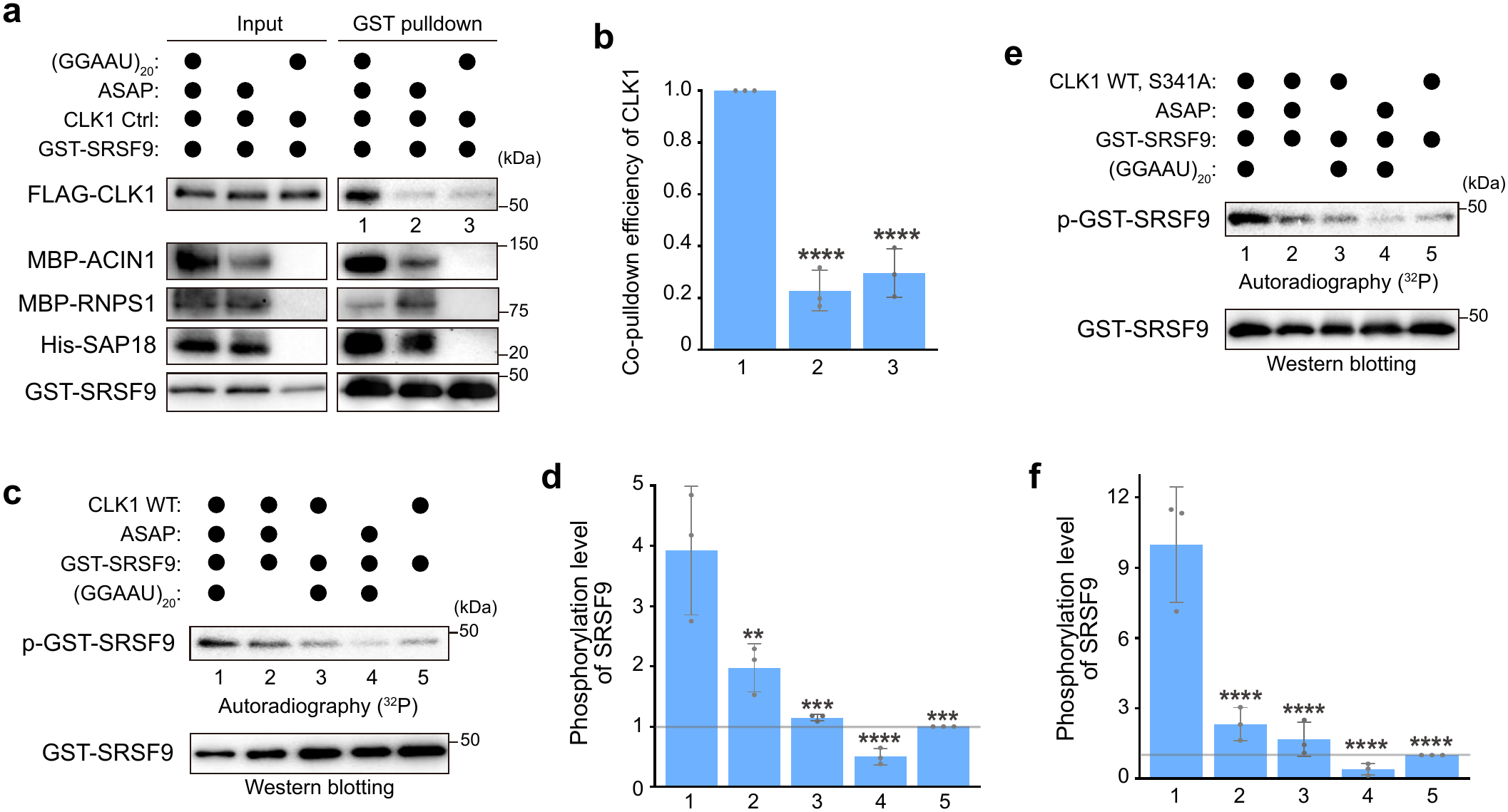
The HSATIII-SRSF-ASAP platform amplifies SRSF phosphorylation through enhanced SRSF-CLK1 association and selective recruitment of activated CLK1. (a, b) In vitro GST pulldown of recombinant GST-SRSF9 with the indicated components (a), and quantification of CLK1 co-pulldown efficiency, calculated as the ratio of CLK1 to SRSF9 (b). Data are shown as mean ± SD. n = 3. Šídák’s multiple-comparisons test was used. **** *P* < 0.0001. (c, d) In vitro SRSF9 phosphorylation by recombinant CLK1 WT in the presence of indicated components, assessed by autoradiography following GST pulldown (c), and quantification of SRSF9 phosphorylation level (d). Data are shown as mean ± SD. n = 3. Šídák’s multiple-comparisons test was used. **** *P* < 0.0001, *** *P* < 0.001, ** *P* < 0.01. (e, f) In vitro SRSF9 phosphorylation by both CLK1 WT and S341A in the same reaction with the indicated components, assessed by autoradiography following GST pulldown (e), and quantification of SRSF9 phosphorylation level (f). Data are shown as mean ± SD. n = 3. Šídák’s multiple-comparisons test was used. **** *P* < 0.0001.

We next examined whether the platform also provides a qualitative advantage by selectively enriching Ser341-phosphorylated CLK1. SRSF9 itself bound CLK1 Ctrl and S341A comparably in vitro (Extended Data Fig. 3h,i) and recruited CLK1 to GGAAU RNA irrespective of its Ser341 phosphorylation status (Extended Data Fig. 6b,c). By contrast, the platform selectively recruited Ser341-phosphorylated CLK1 (Fig. 5i). Because Ser341 phosphorylation enhances CLK1 enzymatic activity toward SRSF9 phosphorylation,^26^ we reasoned that the HSATIII-SRSF-ASAP platform may selectively enrich enzymatically active CLK1 on HSATIII arcRNA and thereby facilitate phosphorylation of SRSFs sequestered in the nSB “reaction crucible”. We therefore reconstituted SRSF9 phosphorylation in the simultaneous presence of Ser341-phosphorylated, catalytically activated, CLK1 WT and the nonphosphorylatable S341A mutant (Extended Data Fig. 6a).^26^ The HSATIII-SRSF-ASAP platform enhanced SRSF9 phosphorylation by more than 10-fold compared with SRSF9 and CLK1 alone (Fig. 6e,f). Omission of either GGAAU or ASAP reduced SRSF9 phosphorylation by approximately 40%. Together with the quantitative enhancement (4-fold) observed with CLK1 WT alone (Fig. 6c,d), these results are consistent with qualitative selection of Ser341-phosphorylated and enzymatically activated CLK1 further promoting efficient substrate phosphorylation. This enhancement was substantially weaker with antisense AUUCC repeat RNAs, which showed substantially weaker SRSF9 binding than GGAAU-repeats (Extended Data Fig. 6d-f), highlighting the importance of sequence-specific RNA recognition by SRSFs for efficient nSB reaction-crucible activity. Thus, the HSATIII-SRSF-ASAP platform enhances SRSF phosphorylation through both quantitative enhancement of enzyme-substrate association and qualitative selection of catalytically activated, Ser341-phosphorylated CLK1. These findings provide the molecular basis for the reaction-crucible activity of nSBs.

### HSATIII co-opts an evolutionarily conserved SRSF-ASAP-CLK1 interaction network

nSBs form in primate cells on HSATIII arcRNAs transcribed from primate-specific satellite III repeats.^13–19^ Our previous work demonstrated that Chinese hamster ovary (CHO) somatic hybrid cells carrying human chromosome 9, which contains a large satellite III repeat region, assembled nSBs using hamster proteins upon thermal stress and accelerated intron detention in hamster pre-mRNAs through the reaction-crucible mechanism during stress recovery.^12^ This finding suggests that HSATIII arcRNA can co-opt pre-existing molecular machinery to establish the reaction crucible. Unlike HSATIII, which is transcribed from primate-specific satellite III repeats, SRSF9 and the ASAP components are highly conserved across mammals, including mouse (Extended Data Fig. 7a-d). We therefore tested whether recombinant mouse Srsf9 and ASAP could assemble the platform on GGAAU-repeat RNA and recruit CLK1 (Extended Data Fig. 7e). Mouse ASAP bound CLK1 comparably to human ASAP (Fig. 7a), and mouse Srsf9 directly interacted with all three ASAP components (Fig. 7b). Moreover, mouse Asap and Srsf9 recruited CLK1 to GGAAU-repeats, whereas recruitment of CLK1 S341A was substantially reduced (Fig. 7c,d), indicating that the mouse Srsf9-ASAP network retains the ability to selectively recruit Ser341-phosphorylated CLK1 to GGAAU-repeats. Strikingly, the platform reconstituted mouse proteins enhanced CLK1-mediated Srsf9 phosphorylation approximately 7-fold compared with Srsf9 and CLK1 alone (Fig. 7e,f). Thus, the SRSF-ASAP-CLK1 network is conserved in mouse and retains the capacity to enhance SRSF phosphorylation on GGAAU-repeat RNA. These findings suggest that the capacity for RNA scaffold-dependent reaction enhancement predates the emergence of primate-specific HSATIII arcRNA. We propose that HSATIII arcRNA co-opted this evolutionarily conserved network to establish a primate-specific reaction crucible that accelerates the normalization of stress-induced splicing during thermal stress recovery (Fig. 7g).

**Figure 7.**
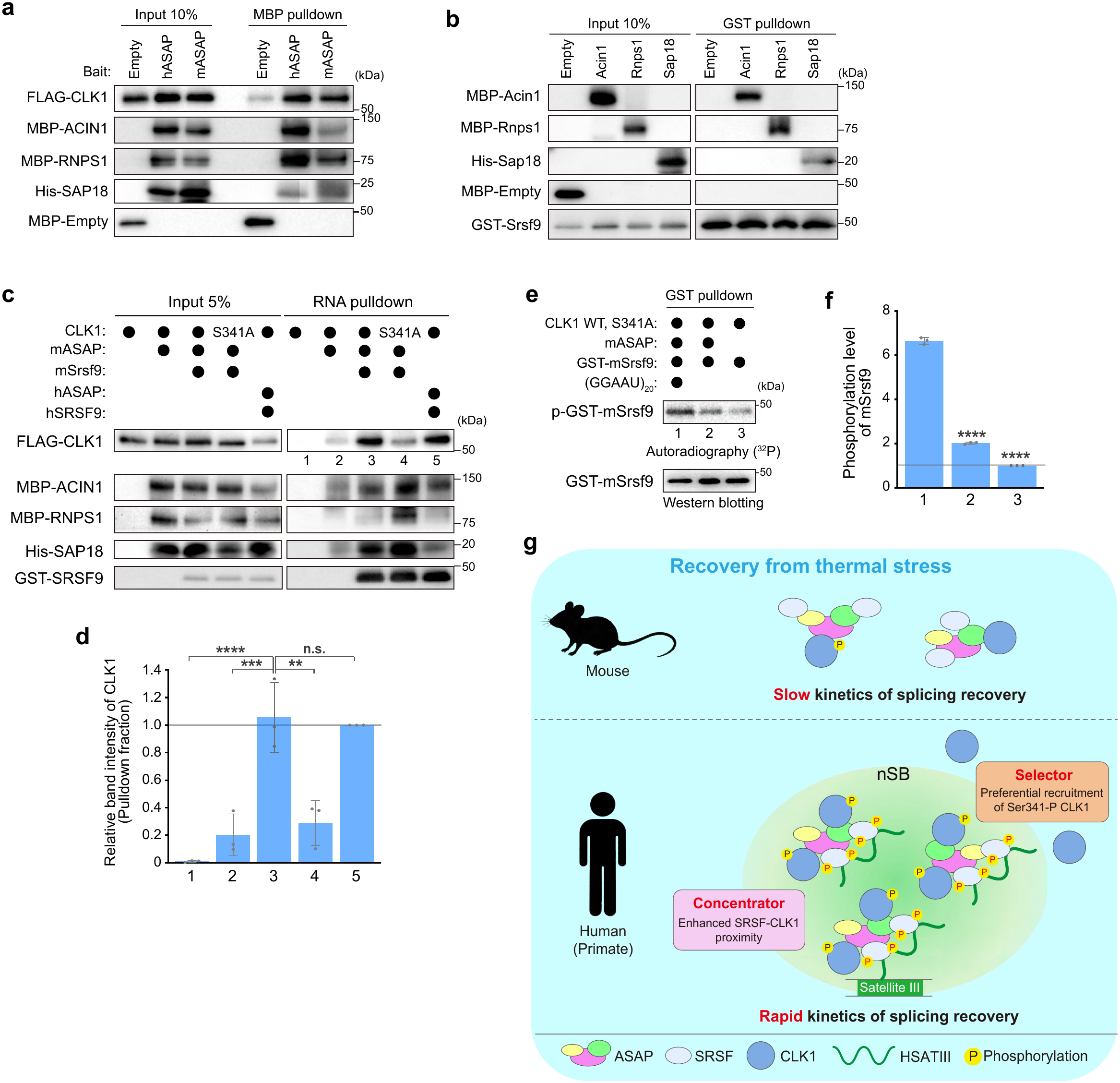
HSATIII co-opts the conserved SRSF-ASAP-CLK1 network to amplify SRSF phosphorylation in an RNA scaffolded reaction crucible. (a) In vitro binding of human ASAP and mouse Asap to human CLK1 using recombinant proteins. (b) In vitro binding of mouse Srsf9 to individual mouse Asap components using recombinant proteins. (c, d) In vitro RNA pulldown using GGAAU-repeat RNA with mouse Srsf9 and Asap (c), and quantification of CLK1 in the pulldown fraction (d). Data are shown as mean ± SD. n = 3. Šídák’s multiple-comparisons test was used. *** *P* < 0.001. ** *P* < 0.01. Human SRSF9 and ASAP were used as positive controls for CLK1 recruitment to GGAAU RNA. (e, f) In vitro mouse Srsf9 phosphorylation by CLK1 WT and S341A in the presence of indicated components, assessed by autoradiography following GST pulldown (e), and quantification of Srsf9 phosphorylation level (f). Data are shown as mean ± SD. n = 3. Šídák’s multiple-comparisons test was used. **** *P* < 0.0001. (g) Schematic model of the evolutionary co-option of a conserved SRSF-ASAP-CLK1 network by primate-specific HSATIII arcRNA. In mice, the conserved protein network lacks an HSATIII RNA scaffold, potentially leading to slower kinetics of SRSF rephosphorylation and splicing recovery following thermal stress. In humans, HSATIII arcRNA organizes the conserved network into nSBs, which function as both molecular concentrators that enhance SRSF-CLK1 association and selectors that preferentially recruit enzymatically activated CLK1, thereby promoting efficient SRSF rephosphorylation and rapid splicing recovery.

## Discussion

Among their various modes of action, MLOs can function as reaction crucibles that spatially and temporally promote specific biochemical reactions. However, how MLOs selectively recruit and organize their functional components required for such reactions remains a central question. nSBs provide a tractable model to address this question because their formation and function can be temporally separated by temperature shifts: they form during thermal stress but begin to function as reaction crucibles specifically during stress recovery. We previously showed that nSBs recruit CLK1 specifically during stress recovery, accelerating SRSF phosphorylation and rapidly reversing stress-induced splicing.^12,18^ This temperature-dependent recruitment requires CLK1 Ser341 phosphorylation,^26^ but how phosphorylated CLK1 is selectively recruited to nSBs remained unknown. Here, we identify the ASAP complex as a phospho-selective recruitment factor that recognizes Ser341-phosphorylated CLK1 and connects it to SRSFs already sequestered on HSATIII arcRNAs. This finding extends the role of ASAP beyond its established function as an EJC-associated accessory complex to include the spatial organization of CLK1-mediated SRSF phosphorylation within nSBs and nSB-mediated splicing control.^27, 29^

ASAP interacts with both SRSFs and CLK1, and HSATIII-bound SRSFs serve as adaptors for ASAP, thereby forming the HSATIII-SRSF-ASAP platform for CLK1 recruitment. Biochemical reconstitution demonstrated that this platform enhances CLK1-SRSF association and promotes SRSF phosphorylation. Importantly, ASAP selectively recognizes Ser341-phosphorylated CLK1, enabling the platform to recruit catalytically activated CLK1, thereby allowing nSBs to function as highly active reaction crucibles.

A key principle emerging from these findings is that nSBs act not only as molecular concentrators but also as selectors of activated enzymes. First, the platform quantitatively enhances CLK1-SRSF association, increasing SRSF9 phosphorylation approximately 4-fold compared with SRSF9 and CLK1 alone (Fig. 6a-d). Second, ASAP selectively recognizes Ser341-phosphorylated CLK1, whose catalytic activity is enhanced by this phosphorylation.^26^ When CLK1 WT and S341A were present together, the platform enhanced SRSF9 phosphorylation by more than 10-fold compared with SRSF9 and CLK1 alone (Fig. 6e,f), consistent with the combined effects of quantitative recruitment and selective enrichment of activated CLK1. Thus, nSBs function as both concentrators and selectors, maximizing reaction-crucible output through efficient enzyme-substrate association and preferential recruitment of activated enzyme (Fig. 7g). Moreover, the recovery-specific phosphorylation of CLK1 at Ser341 enables nSBs to function as reaction crucibles specifically during stress recovery.

This dual activity is supported by the multivalent organization of the platform. Multiple SRSF9 molecules can bind to a single HSATIII RNA through its repetitive GGAAU sequences. SRSF9 can independently interact with each of the three ASAP components, while CLK1 can interact with two of them, ACIN1 and RNPS1. These multiple RNA-protein and protein-protein interactions within the platform can efficiently bring the kinase and its substrate into close proximity (Fig. 5k). Accordingly, neither the RNA scaffold nor ASAP alone is sufficient for maximal reaction enhancement. Moreover, replacing GGAAU-repeats with antisense AUUCC-repeats substantially weakens SRSF9 phosphorylation (Extended Data Fig. 6d,e), indicating that RNA sequence is critical for reaction-crucible activity of nSBs. SRSFs therefore serve a dual role as both substrates of CLK1 and adaptors that couple the ASAP-CLK1 machinery to HSATIII. These findings illustrate how an arcRNA can organize proteins that do not themselves efficiently bind the arcRNA and how multivalent interactions can convert molecular recruitment into productive biochemical activity.

The phospho-selective nature of this system also provides a mechanism for temporal control of nSB reaction-crucible function. Because RIOK2 phosphorylates CLK1 at Ser341 specifically during stress recovery, phosphorylated CLK1 is selectively recruited to the platform during this phase. RIOK2 phosphorylates CLK1 in the nucleoplasm rather than within nSBs,^26^ suggesting that CLK1 phosphorylation and its recruitment to the platform are spatially separated. Interestingly, ASAP shows only modest enrichment in nSBs despite its critical role in CLK1 recruitment (Fig. 1d–f and Extended Data Fig. 1f– h), indicating that pronounced accumulation of ASAP is not required for its recruitment function. How Ser341 phosphorylation promotes ASAP binding and CLK1 kinase activity remains unclear. Given that phosphorylation can alter the accessibility of protein-interaction surfaces,^39–41^ Ser341 phosphorylation may remodel CLK1 conformation, thereby promoting kinase activation and interaction with ASAP.

Finally, our findings suggest a mechanism by which evolutionarily novel lncRNAs can acquire beneficial biochemical functions (Fig. 7). SRSF9 contains a canonical RRM1 and a pseudo-RRM2, with RRM2 recognizing AGGA-containing sequences through a conserved noncanonical RNA-binding mode.^42,43^ The conservation of this RNA-recognition module, together with conserved SRSF-ASAP-CLK1 interactions and RNA scaffold-dependent enhancement of CLK1-mediated SRSF phosphorylation observed with mouse proteins, suggests that the molecular networks exploited by HSATIII arcRNA predate the emergence of satellite III repeat regions in primate genomes. This model is consistent with our previous reconstitution of nSB reaction-crucible activity in splicing regulation in CHO somatic hybrid cells carrying human chromosome 9, from which HSATIII is transcribed.^12^ This finding demonstrates that pre-existing trans-acting machinery can support and shape HSATIII function. Together, our findings reveal how HSATIII arcRNAs co-opt conserved protein interaction networks to assemble multivalent platforms that selectively recruit activated enzymes and establish functional reaction crucibles (Fig. 7g). Because lncRNAs generally show limited sequence conservation and often lineage-specific transcription units,^44–46^ evolutionarily novel arcRNAs may similarly acquire function by co-opting pre-existing RNA-recognition and protein-interaction networks rather than evolving numerous new molecular interactions. Such co-option may enable rapidly evolving lncRNAs to build new biochemical compartments.

## Methods

### Cell culture

HeLa,^12^ Flp-In™ T-REx™293 (Thermo Fisher Scientific, R78007), and HEK 293 (ATCC, CRL-1573) cells were maintained in DMEM supplemented with 10% fetal calf serum and antibiotics at 37°C under 5% CO_2_, as described previously.^26^

### Thermal stress and chemical treatment

Thermal stress was induced by incubating cells at 42℃ under 5% CO_2_. Chemical treatments were performed by replacing the culture medium with pre-warmed medium containing the indicated compounds.

### Plasmid construction and transfection

Human cDNA fragments were amplified by PCR and cloned into tag-containing pcDNA5/FRT/TO/i vectors^4^ or pcDNA3 vectors for expression in cultured cells. For recombinant protein production in E. coli, the corresponding fragments were cloned into pET28a or pGEX-6P-1.^47^ Deletion constructs, amino acid substitutions, and siRNA-resistant variants were generated by inverse PCR or In-Fusion Snap Assembly (TaKaRa). Plasmids used in this study are listed in Supplementary Table 2, and primer sequences are provided in Supplementary Table 3.

### Plasmid transfection and knockdown using siRNA

Plasmid transfection and siRNA-mediated knockdown were performed as described previously.^26^ Briefly, plasmids were transfected using TransIT^®^-LT1 Transfection Reagent. siRNAs were transfected using Lipofectamine RNAiMAX (Thermo Fisher Scientific) 48 h before analysis, and plasmids were transfected 24 h before analysis. siRNAs were transfected at 20 nM (10 nM each for double knockdown). Plasmids and siRNAs used in this study are listed in Supplementary Table 2 and 5, respectively.

### Co-immunoprecipitation

Co-immunoprecipitation was performed as described previously.^26^ Briefly, cell lysates were incubated with antibody-conjugated Dynabeads^TM^ Protein G (Invitrogen) at 4°C. After washing, bound proteins were eluted with SDS sample buffer or FLAG peptide and analyzed by western blotting.

### Mass spectrometry

Mass spectrometry was performed as described previously.^26^ Briefly, proteins recovered by immunoprecipitation were precipitated, reduced, alkylated, and sequentially digested with lysyl endopeptidase and trypsin. The resulting peptides were analyzed by LC-MS/MS using an Evosep One LC system coupled to a Q-Exactive HF-X mass spectrometer (Thermo Fisher Scientific). Protein identification was performed using Proteome Discoverer with the SEQUEST search engine.

### RNA-FISH and immunofluorescence

RNA-FISH and immunofluorescence were performed as described previously.^26^ HSATIII and the MS2 RNA used in Figure 4 and Extended Data Figure 4 were detected using fluorescently labeled antisense oligonucleotides, and immunofluorescence was performed using the indicated antibodies. For the MS2 RNA FISH, Blocking Reagent (Roche) was used for blocking and antibody incubation during subsequent immunofluorescence. Images were acquired using an LSM900 confocal microscope (Zeiss) and analyzed with ImageJ (FIJI). Antibodies and oligonucleotides are listed in Supplementary Table 4 and 5, respectively.

### HSATIII ChIRP

HSATIII ChIRP was performed as described previously.^12,22,26^ Briefly, PFA-crosslinked cells were lysed and sonicated, and HSATIII-containing complexes were hybridized with a biotinylated HSATIII antisense oligonucleotide and recovered using Dynabeads™ MyOne™ Streptavidin C1 (Invitrogen). Bound proteins were eluted and analyzed by western blotting, whereas recovered RNA was purified after reversal of crosslinking.

### Western blotting

Western blotting was performed as described previously.^26^ Protein signals were detected using a ChemiDoc Touch imaging system (Bio-Rad) and quantified with ImageJ (FIJI). Antibodies used in this study are listed in Supplementary Table 4.

### RNA extraction and RT-qPCR

RNA extraction and quantitative RT-PCR were performed as described previously.^26^ Briefly, total RNA was extracted from cultured cells or ChIRP samples, treated with DNase, and reverse-transcribed into cDNA. Quantitative PCR was performed using KAPA SYBR Fast qPCR Kit (Nippon Genetics) on a LightCycler 480 System II (Roche). Primer sequences are provided in Supplementary Table 3.

### Expression and purification of recombinant proteins

Recombinant proteins were expressed and purified from E. coli essentially as described previously.^26^ Briefly, His-tagged proteins were expressed in Rosetta-gami 2 (DE3) cells and purified using Ni-NTA agarose (Wako). Where indicated, GST- or MBP-tagged proteins were further purified using Glutathione Sepharose 4 Fast Flow (Cytiva) or Amylose Resin (NEB), respectively. Purified proteins were dialyzed against buffer containing 20 mM Tris-HCl (pH 7.5), 1 mM DTT, 0.2 mM EDTA, and 10% glycerol, supplemented with 150 mM NaCl. Recombinant ACIN1 and RNPS1 were instead dialyzed against the same buffer containing 500 mM NaCl. GST-SRSF9 WT and mutant proteins were recovered from the insoluble fraction by solubilization in 20 mM Tris-HCl (pH 7.5), 500 mM NaCl, and 6 M urea, followed by purification using Ni-NTA agarose (Wako). Purified proteins were subsequently dialyzed against buffer containing 20 mM Tris-HCl (pH 7.5), 500 mM NaCl, 1 mM DTT, 0.2 mM EDTA, and 10% glycerol.

For purification from mammalian cells, FLAG-CLK1 was expressed in Flp-In™ T-REx™ 293 cells by induction with 1 μg/mL doxycycline for 24 h. Cells were lysed in IP-1000 buffer (20 mM Tris-HCl, pH 7.5, 1 M NaCl, and 0.5% NP-40) supplemented with cOmplete Protease Inhibitor Cocktail (Sigma), PhosSTOP (Roche), and 1 mM PMSF. After sonication and centrifugation, clarified lysates were incubated with anti-FLAG antibody-conjugated Dynabeads^TM^ Protein G (Invitrogen) at 4°C for 3 h. Beads were washed five times with IP-1000 buffer supplemented with PhosSTOP, and FLAG-CLK1 was eluted with 3× FLAG peptide (Protein Ark).

MBP-tagged ACIN1-S WT, ACIN1-S ΔRSB, RNPS1 WT, and RNPS1 ΔRRM were expressed in HEK293 cells by transfection with PEI MAX (Cosmo Bio). Cells were lysed in IP-1000 buffer supplemented with cOmplete Protease Inhibitor Cocktail (Sigma), PhosSTOP (Roche), and 1 mM PMSF. After sonication and centrifugation, clarified lysates were incubated with Amylose Resin (NEB) at 4°C. The resin was washed with IP-1000 buffer, and bound proteins were eluted with maltose-containing buffer (20 mM Tris-HCl (pH 7.5), 500 mM NaCl, 50 mM maltose, 1 mM DTT, 0.2 mM EDTA, 10% glycerol). Purified proteins were subsequently dialyzed against the same buffer without maltose.

### In vitro binding assay

In vitro binding assays were performed essentially as described previously.^26^ Purified bait and target proteins were mixed at a final NaCl concentration of 150 mM and incubated on ice for 30 min. Protein complexes were then captured using Amylose Resin (NEB) for MBP-tagged bait proteins or Glutathione Sepharose 4 Fast Flow (Cytiva) for GST-tagged bait proteins. For binding assays between His-SAP18 and CLK1, protein complexes were captured using anti-His antibody-conjugated Dynabeads Protein G (Invitrogen). After incubation at 4℃ for 1 h, the resins or beads were washed five times, and bound proteins were eluted with SDS sample buffer and analyzed by SDS-PAGE and western blotting.

For binding assays using the ASAP complex, recombinant ASAP components were first purified using Ni-NTA agarose (Wako) and eluted with imidazole. The eluted proteins were subsequently purified using Amylose Resin (NEB) to isolate the assembled ASAP complex. The bead-bound ASAP complex was then incubated with the indicated target proteins under the same binding conditions described above. After washing, bound proteins were eluted with SDS sample buffer and analyzed by SDS-PAGE and western blotting.

### In vitro kinase assay

To prepare RIOK2-phosphorylated CLK1 for subsequent in vitro assays, Glutathione Sepharose 4B (Cytiva)-bound RIOK2 was mixed with recombinant CLK1 purified from E. coli and incubated at 37℃ with shaking at 800 rpm for 1 h in IP-150 buffer (20 mM Tris-HCl, pH 7.5, 150 mM NaCl, and 0.5% NP-40) supplemented with 10 mM MgCl_2_, 2 mM DTT, and 1 mM ATP. Following the kinase reaction, the NaCl concentration was increased to 500 mM by addition of 5 M NaCl, and the mixture was rotated at 4℃ for 1 h. IP-1000 buffer was then added, and the supernatant containing phosphorylated CLK1 was recovered after removal of the RIOK2-bound beads. The NaCl concentration was subsequently adjusted to 150 mM by addition of IP-0 buffer (20 mM Tris-HCl, pH 7.5, and 0.5% NP-40), and phosphorylated CLK1 was concentrated using an Amicon centrifugal filter unit (Merck).

RIOK2-mediated phosphorylation of CLK1 was verified by an in vitro kinase assay using bead-bound RIOK2 and [γ-³²P] ATP (Revvity), as described previously ^26^.

### MS2-tethering assay

The MS2-MCP tethering assay was performed based on previously described approaches.^4,48,49^ The RNA containing 18× MS2- and 24× PP7-binding sites was stably expressed in Flp-In™ T-REx™293 cells. Cells were seeded onto poly-L-lysine-coated coverslips and, 24 h later, transfected with plasmids expressing the indicated MCP-fused proteins and CLK1. Expression of the scaffold RNA was induced with 1 μg /mL doxycycline at the time of transfection. After 24 h, cells were fixed and subjected to RNA-FISH and immunofluorescence. The MS2 RNA was detected using a Cy5-labeled oligonucleotide probe complementary to a linker sequence within the 24x PP7 array.

### In vitro transcription

(GGAAU)_20_ repeat RNA was generated by in vitro transcription using an annealed DNA template containing a T7 promoter and the (ATTCC)_20_ repeat sequence. Biotinylated (GGAAU)_20_ and (AUUCC)_20_, and unlabeled (GGAAU)_20_ repeat RNAs were synthesized using the MEGAscript™ T7 Transcription Kit (Thermo Fisher Scientific), whereas FITC-labeled (GGAAU)_20_ repeat RNA was synthesized using the RNA Labeling Kit (Roche), according to the manufacturers’ instructions. Following treatment with RQ1 DNase (Promega), the transcribed RNAs were purified using the RNA Clean & Concentrator-5 kit (Zymo Research) or by phenol-chloroform extraction followed by ethanol precipitation. Template DNA sequences are listed in Supplementary Table 3.

### In vitro RNA pulldown

Biotinylated (GGAAU)_20_ repeat RNA was heated at 90℃ for 2 min and immediately placed on ice for 2 min. RNA was then refolded by addition of an equal volume of 2× RNA structure buffer (20 mM Tris-HCl, pH 7.4, 200 mM KCl, and 20 mM MgCl₂). Refolded RNA was immobilized on Dynabeads MyOne Streptavidin C1 (Thermo Fisher Scientific) by rotation at 4℃ for 1 h.

Recombinant proteins were diluted in IP-150 buffer supplemented with cOmplete Protease Inhibitor Cocktail (Sigma), PhosSTOP (Roche), 1 mM PMSF, and RNase inhibitor (Thermo Fisher Scientific), and precleared with Dynabeads MyOne Streptavidin C1 at 4℃ for 1 h. Precleared proteins were then incubated with RNA-conjugated beads at 4℃ for 1 h with rotation. The beads were washed five times with IP-150 buffer, and bound proteins were eluted with SDS sample buffer and analyzed by SDS-PAGE and western blotting.

For the reconstitution assay shown in Figure 5i, the recombinant ASAP components were first mixed and incubated on ice for 30 min. SRSF9 and CLK1 were then added to the preassembled ASAP complex, and the resulting protein mixture was subjected to RNA pulldown as described above.

### Electrophoretic mobility shift assay (EMSA)

FITC-labeled (GGAAU)_20_ repeat RNA was incubated with recombinant proteins in EMSA buffer containing 250 mM sucrose, 2 mM MgCl_2_, 20 mM Tris-HCl (pH 7.5), and RNase inhibitor (Thermo Fisher Scientific) at 30°C for 15 min. Samples were subsequently resolved by native 6% PAGE. FITC-labeled RNAs were detected using a ChemiDoc Touch imaging system (Bio-Rad) and quantified with ImageJ (FIJI). The percentage of shifted RNA was calculated relative to the total RNA signal in each lane, with signals retained in the wells included in the shifted RNA fraction.

### In vitro reconstitution of SRSF9 phosphorylation

In vitro-transcribed and refolded (GGAAU)_20_ repeat RNA was pre-incubated with SRSF9, while ASAP components were separately pre-incubated with CLK1 WT and S341A at room temperature for 15 min. The two pre-incubated mixtures were then combined in reaction buffer containing 5 mM MgCl_2_, 2 mM DTT, and 10 μM unlabeled ATP, with a final NaCl concentration of 150 mM. Phosphorylation was initiated by the addition of [γ-^32^P] ATP (Revvity), and the reactions were incubated at 37℃ for 15 min. GST-SRSF9 was then recovered using Glutathione Sepharose 4 Fast Flow (Cytiva) in IP-500 buffer by rotation at 4℃ for 1 h. The beads were washed five times with IP-500 buffer, and bound proteins were denatured in SDS sample buffer and analyzed by SDS-PAGE and autoradiography.

### Statistics and reproducibility

RT-qPCR data were processed using LightCycler 480 software (version 1.5.1; Roche). Western blotting, RNA-FISH and immunofluorescence images were quantified using ImageJ (FIJI), with Pearson’s correlation coefficients determined using the Coloc 2 plugin. Statistical analyses were performed using GraphPad Prism 10.

All experiments were performed at least twice independently, with similar results obtained. No statistical method was used to predetermine sample size, and no data were excluded from the analyses. Data are presented as mean ± S.D. where indicated. For imaging analyses, n represents individual cells analyzed within a single representative experiment; the corresponding experiments were independently repeated at least twice with similar results. For biochemical assays and RT-qPCR analyses, n represents independent experiments unless otherwise stated.

Kruskal-Wallis test followed by Dunn’s multiple-comparisons test was used for Fig. 2b,d,f,h,j and Extended Data Figs. 2f,m,o and 3d,f. Mann-Whitney U test was used for Fig. 4f,i and Extended Data Figs. 2i, 4e,g and 5i. Two-tailed t-test was used for Figs. 1a and 3j,m and Extended Data Figs. 3i and 6c. One-way ANOVA followed by Šídák’s multiple-comparisons test was used for Figs. 2k, 3b,e,h,o, 5d,h,j, 6b,d,f and 7d,f and Extended Data Figs. 2q, 5l and 6e. Statistical significance is denoted as follows: * 0.01 < *P* < 0.05, ** 0.001 < *P* < 0.01, *** 0.0001 < *P* < 0.001 and **** *P* < 0.0001. Exact n values are provided in the corresponding figure legends.

## Data availability

All data are available in the main text or the supplementary materials. The mass spectrometry proteomics data reanalysed in this study were derived from the same raw datasets reported in our previous study^26^ and deposited in the Japan ProteOme Standard Repository (jPOST) under accession numbers JPST004102 and JPST004490.

## Acknowledgments

We thank the FBS Core Facility at the University of Osaka for technical assistance. We thank the members of the Hirose laboratory for valuable discussions, especially Ichiro Taniguchi and Naoko Fujiwara for guidance on in vitro analyses. We also thank Akila Mayeda for valuable discussions.

## Author contributions

T.U., K.N. and T.H. conceived and designed the study. T.U. performed most experiments and analyses. T.Y. set up the MS2-tethering assays used in Fig. 4 and Extended Data Fig. 4. S.A. performed and analyzed the mass spectrometry experiments. T.U. prepared the figures. K.N. and T.H. supervised the study. T.H., S.A., T.Y. and K.N. acquired funding. T.U., K.N. and T.H. wrote the original manuscript, and all authors reviewed and edited the manuscript.

---

**Extended Data Fig. 1.**
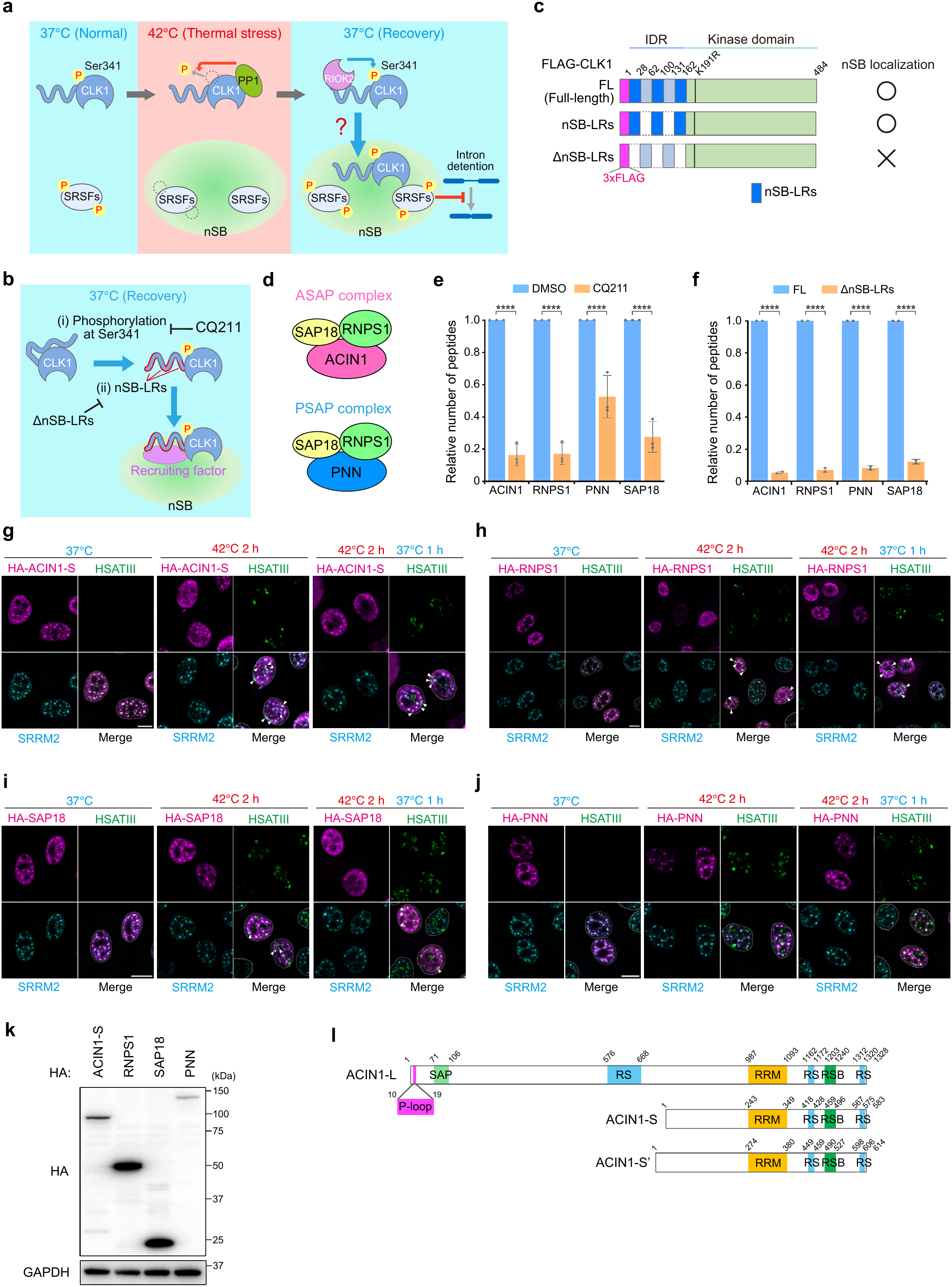
Identification and localization of candidate CLK1 recruitment factors. (a) Schematic of the temperature-dependent regulation of CLK1 Ser341 phosphorylation and its localization to nSBs, thereby regulating intron detention during recovery. During thermal stress, PP1-mediated dephosphorylation of CLK1 Ser341 prevents CLK1 recruitment to nSBs, whereas RIOK2-mediated Ser341 phosphorylation during recovery promotes CLK1 localization to nSBs and nSB-mediated splicing regulation. However, how phosphorylated CLK1 is recruited to nSBs remains unclear. (b) Schematic of the strategy used to identify candidate recruitment factors for CLK1 localization to nSBs. Candidate factors were expected to preferentially associate with Ser341-phosphorylated CLK1 during recovery, as assessed by RIOK2 inhibition with CQ211, and to interact with CLK1 through the N-terminal nSB-LRs, as assessed using the ΔnSB-LRs mutant. nSB-LRs are outlined in red. (c) Schematic of CLK1 constructs. Full-length (FL) CLK1 and mutants lacking nSB-localization regions (nSB-LRs) within the N-terminal IDR are shown, together with their nSB localization. (d) Schematic representation of the ASAP and PSAP complexes. ASAP consists of ACIN1, RNPS1, and SAP18, whereas PSAP consists of PNN, RNPS1, and SAP18. (e, f) Peptide counts of the indicated ASAP and PSAP components identified by CLK1 coIP-MS in the presence or absence of the RIOK2 inhibitor CQ211 (e) or with full-length CLK1 or nSB-LR deletion mutants (f). Data are shown as mean ± SD. n = 3 (e); n = 2 (f). (g-j) Localization of HA-tagged ACIN1, RNPS1, SAP18, and PNN, respectively, to HSATIII-scaffolded nSBs and SRRM2-marked nuclear speckles under the indicated temperature conditions. Nuclei were stained with DAPI. Scale bars, 10 μm. Arrowheads indicate colocalization with nSBs. (k) Expression of HA-tagged ASAP and PSAP components validated by Western blot. (l) Schematic representation of ACIN1 protein isoforms. P-loop, phosphate-binding loop; SAP, SAF-A/B, Acinus and PIAS domain; RS, serine-arginine-rich region; RRM, RNA recognition motif; RSB, RNPS1-SAP18 binding domain.

**Extended Data Fig. 2.**
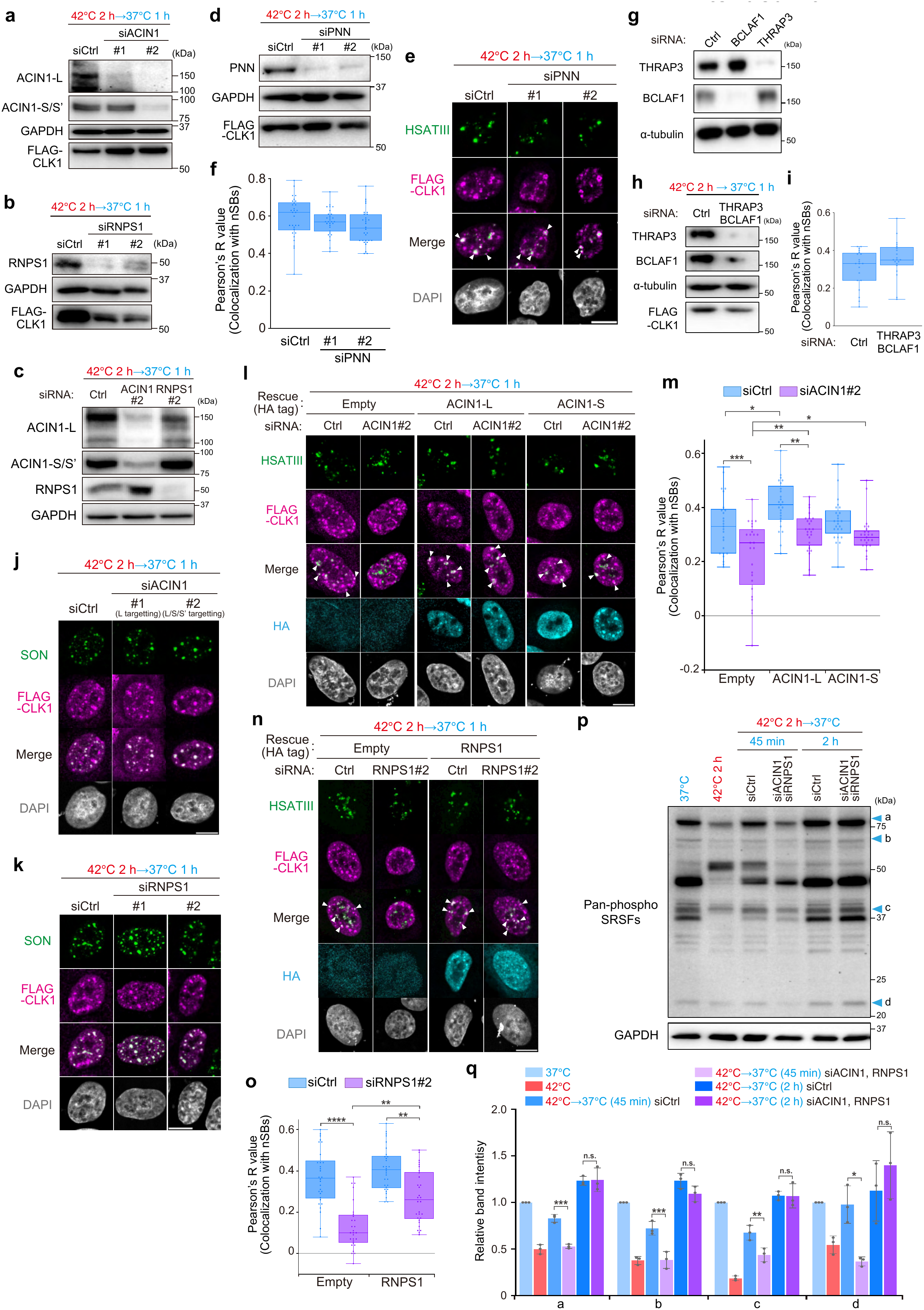
The ASAP complex is required for the reaction-crucible function of nSBs. (a, b, d, g, h) Validation of ACIN1 (a), RNPS1 (b), PNN (d), and individual (g) or combined (h) BCLAF1 and THRAP3 knockdown by Western blot. ACIN1 was depleted using an ACIN1-L-specific siRNA (#1) or an siRNA targeting all ACIN1 isoforms (#2). GAPDH and α-tubulin were used as internal controls. (c) Western blot analysis of RNPS1 and ACIN1 expression following ACIN1 and RNPS1 knockdown, respectively. (e, f) CLK1 localization to nSBs following PNN knockdown (e) and quantification of CLK1-HSATIII colocalization using Pearson’s correlation coefficients (R) within nuclei (f). Nuclei were stained with DAPI. Scale bars, 10 μm. n = 30. Arrowheads indicate CLK1 colocalization with nSBs. Kruskal-Wallis test followed by Dunn’s multiple-comparisons test was used. No statistically significant difference was observed. (i) Quantification of CLK1-HSATIII colocalization following combined BCLAF1 and THRAP3 knockdown using Pearson’s correlation coefficients (R) within nuclei. n = 30. Mann-Whitney U test was used. No statistically significant difference was observed. (j, k) CLK1 localization to nuclear speckles following ACIN1 (j) or RNPS1 (k) knockdown. (l, n) Rescue of CLK1 localization to nSBs by expression of siRNA-resistant ACIN1 following ACIN1 knockdown (l) and siRNA-resistant RNPS1 following RNPS1 knockdown (n). (m, o) Quantification of CLK1-HSATIII colocalization in (l) and (n) using Pearson’s correlation coefficients (R) within nuclei. n = 30. Kruskal-Wallis test followed by Kruskal-Wallis test followed by Dunn’s multiple-comparisons test was used. **** *P* < 0.0001, *** *P* < 0.001, ** *P* < 0.01, * *P* < 0.05. (p, q) SRSF phosphorylation under basal conditions, thermal stress, early recovery (45 min) and late recovery (2 h) following combined ACIN1 and RNPS1 knockdown, assessed by Western blot using the pan-phospho-SRSF antibody 1H4 (p), and quantification of band intensities (q). GAPDH was used as a loading control. Band intensities were normalized to GAPDH. Šídák’s multiple-comparisons test was used. ** *P* < 0.01, * *P* < 0.05. Data are shown as mean ± SD. n = 3.

**Extended Data Fig. 3.**
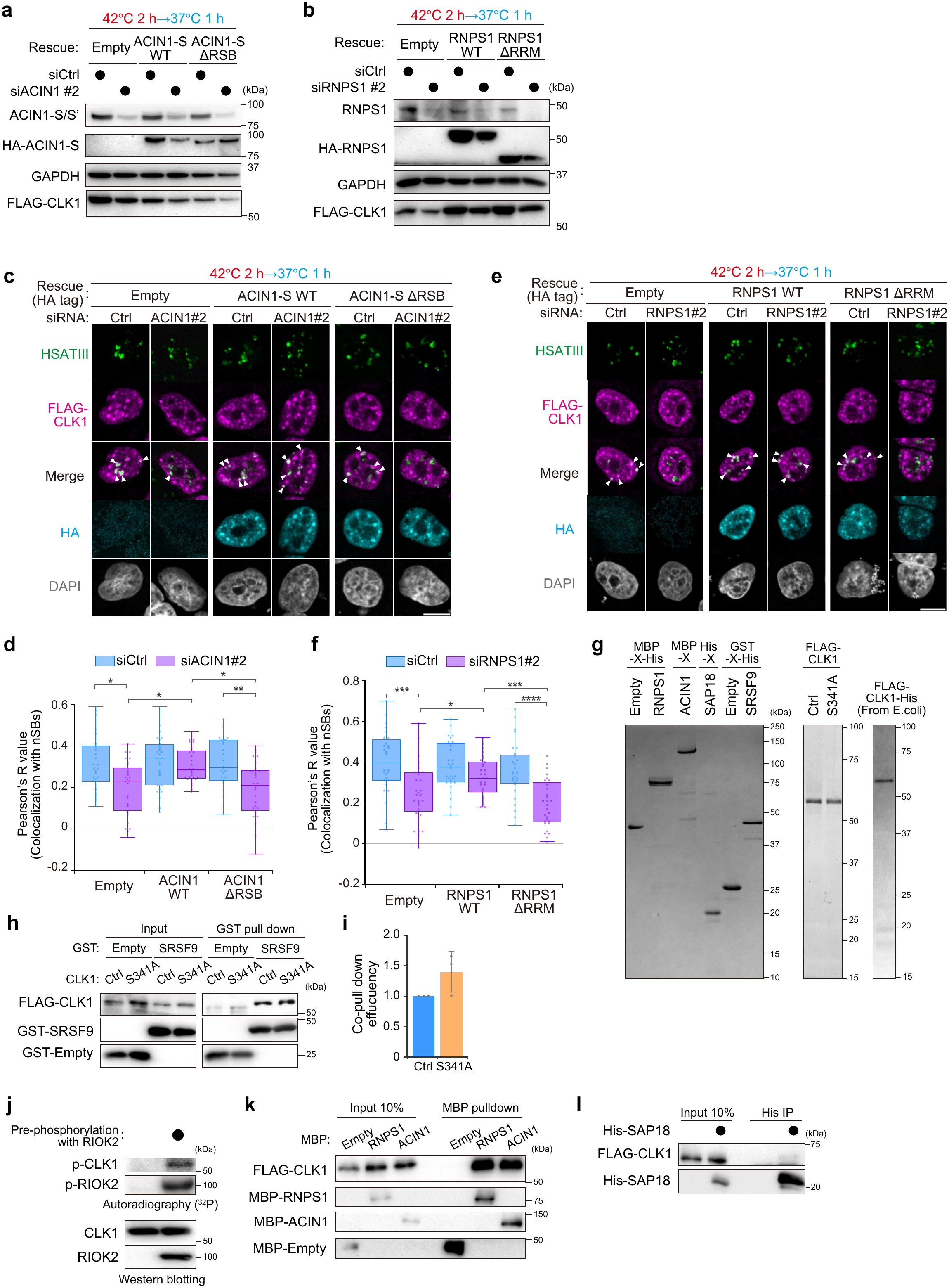
Characterization of the interaction between the ASAP complex and CLK1. (a, b) Western blot analysis validating rescue with siRNA-resistant ACIN1 WT or ΔRSB following ACIN1 knockdown (a), and siRNA-resistant RNPS1 WT or ΔRRM following RNPS1 knockdown (b). (c, e) CLK1 localization to nSBs following ACIN1 (c) or RNPS1 (e) knockdown and rescue with the indicated siRNA-resistant constructs. Nuclei were stained with DAPI. Scale bars, 10 μm. Arrowheads indicate CLK1 colocalization with nSBs. (d, f) Quantification of CLK1-HSATIII colocalization in (c) and (e) using Pearson’s correlation coefficients (R) within nuclei. n = 30. Kruskal-Wallis test followed by Dunn’s multiple-comparisons test was used. **** *P* < 0.0001, *** *P* < 0.001, ** *P* < 0.01, * *P* < 0.05. (g) SDS-PAGE and CBB staining or SYPRO Ruby^®^ staining of purified recombinant proteins used for in vitro binding assays. (h, i) In vitro binding of SRSF9 to control (Ctrl) and S341A CLK1 recombinant proteins (h) and quantification of co-pulldown efficiency (i), calculated as the ratio of CLK1 to SRSF9. Data are shown as mean ± SD. n = 3. Two-tailed t test was used. No statistically significant difference was observed. (j) Validation of CLK1 Ser341 phosphorylation following in vitro phosphorylation by immobilized RIOK2, assessed by autoradiography and Western blot. (k, l) In vitro binding assays examining the interactions of CLK1 with MBP-ACIN1 and MBP-RNPS1 (k) and His-SAP18 (l) using recombinant proteins.

**Extended Data Fig. 4.**
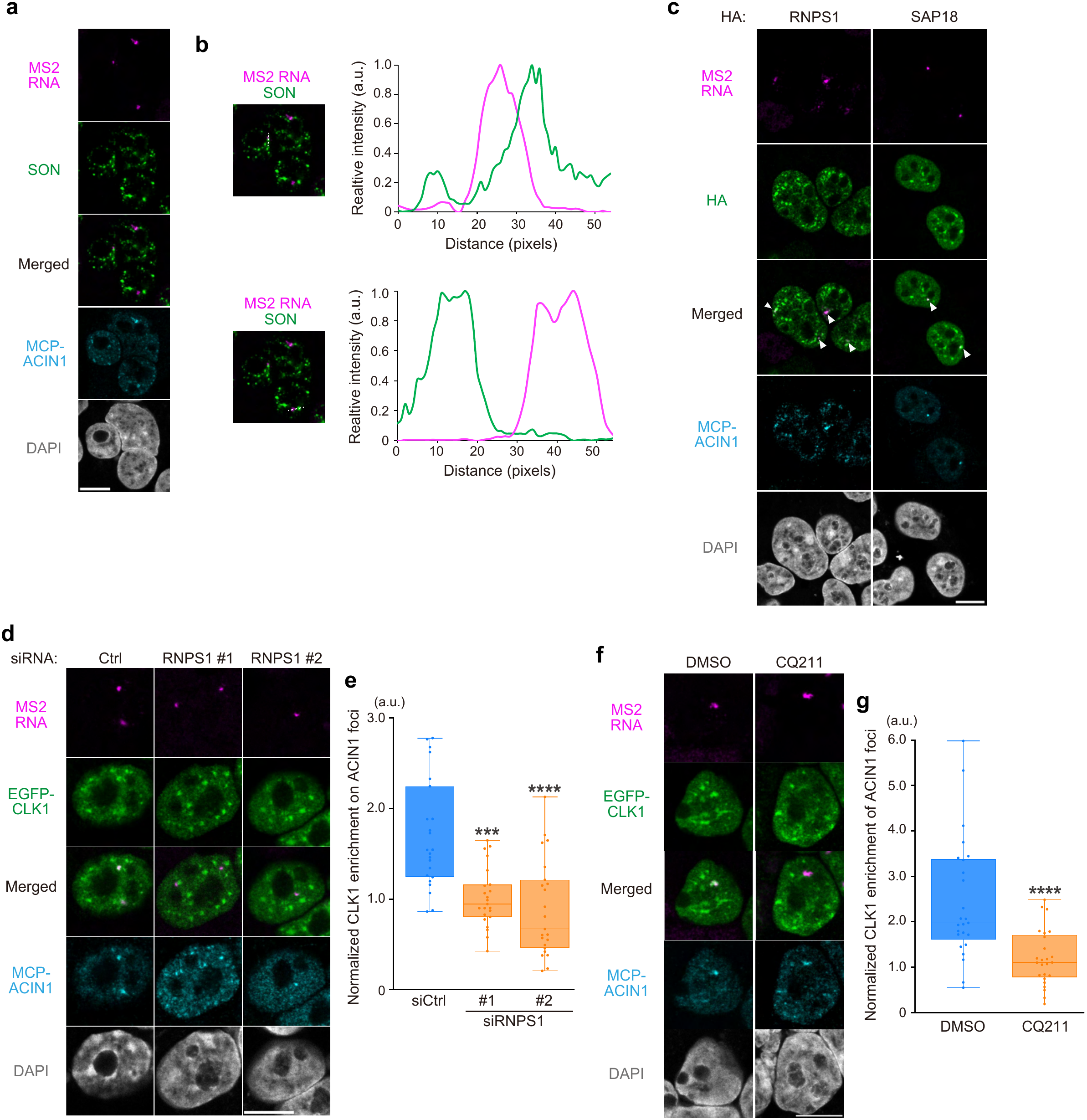
Characterization of ASAP-mediated CLK1 recruitment in the MS2 tethering assay. (a, b) Spatial relationship between MCP-ACIN1-tethered MS2 RNA foci and SON-marked nuclear speckles (a), with line-scan profiles of MS2 RNA and SON fluorescence along the indicated lines (b). Nuclei were stained with DAPI. Scale bars, 10 μm. (c) Recruitment of HA-tagged RNPS1 and SAP18 to MS2 RNA foci tethered with MCP-ACIN1. White arrowheads indicate colocalization of MS2 RNA foci with HA-RNPS1 or HA-SAP18. (d, f) Localization of EGFP-CLK1 to MCP-ACIN1-tethered MS2 RNA foci following RNPS1 knockdown (d) or in the presence or absence of CQ211 (f). siRNA, 20 nM (d). Cells were treated with DMSO or CQ211 (10 μM) for 3 h (f). (e, g) Quantification of CLK1 recruitment in (d) and (f), respectively. CLK1 intensity within MS2 RNA foci was normalized to nuclear CLK1 intensity and MS2 RNA focus intensity. n = 23 (e); n = 26 (g). Mann-Whitney U test was used. **** *P* < 0.0001, *** *P* < 0.001.

**Extended Data Fig. 5.**
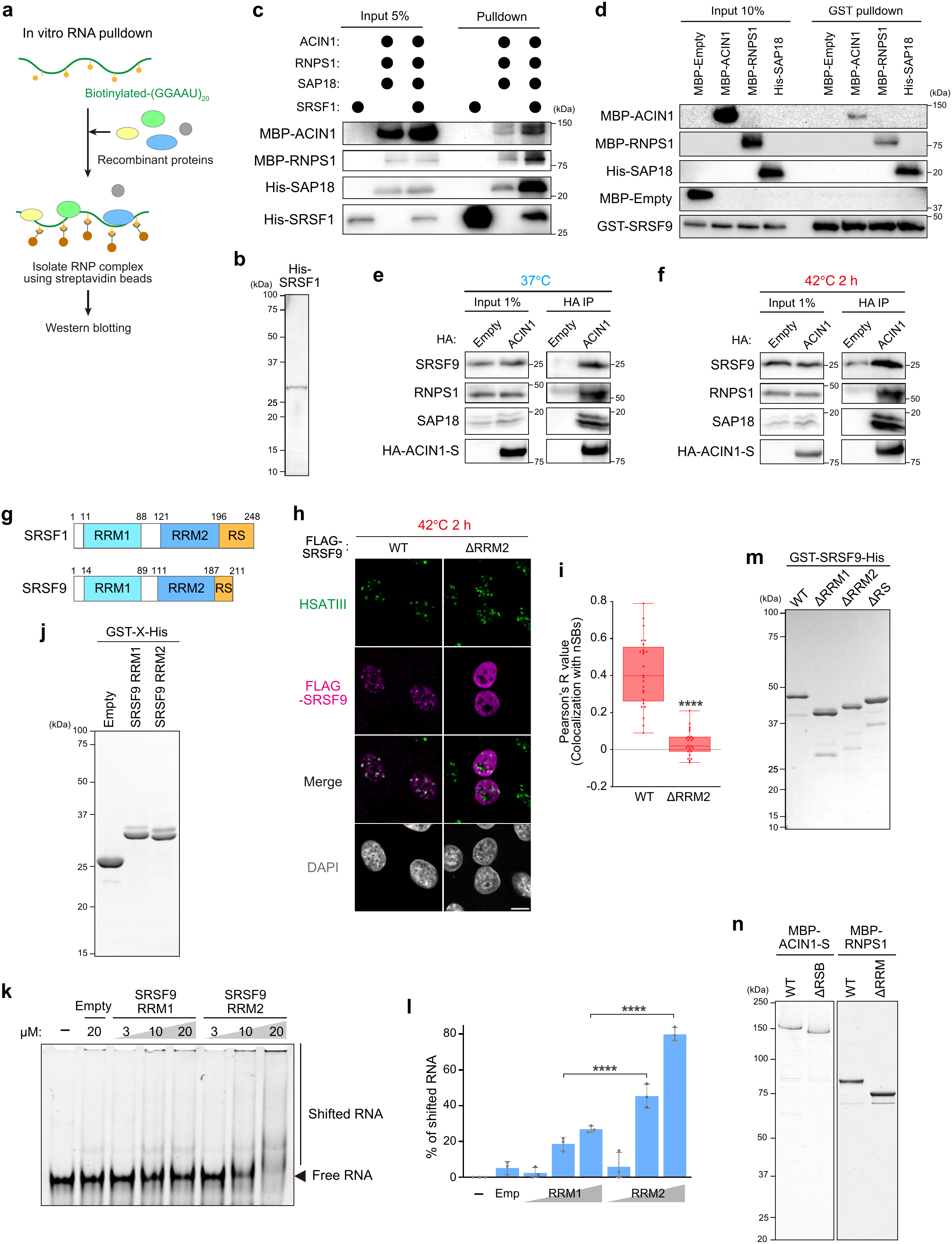
Characterization of the HSATIII-SRSF-ASAP platform. (a) Schematic of the in vitro RNA pulldown assay using recombinant proteins and biotinylated GGAAU-repeat RNA designed to mimic HSATIII arcRNAs. RNA-protein complexes were recovered using streptavidin beads. (b, j, m, n) SDS-PAGE and CBB or SYPRO Ruby^®^ staining of recombinant SRSF1 (b), isolated SRSF9 RRM1 and RRM2 domains (j), SRSF9 deletion mutants (m), and ACIN1 ΔRSB and RNPS1 ΔRRM (n). (c) In vitro RNA pulldown assay using biotinylated GGAAU-repeat and recombinant SRSF1, ASAP, or their combination. (d) In vitro GST pulldown using GST-SRSF9 as bait to assess binding to individual ASAP components using their recombinant proteins. (e, f) Co-IP of HA-ACIN1 to evaluate ASAP-SRSF9 interaction under normal temperature conditions (37°C) (e) and thermal stress (42°C 2 h) (f). (g) Schematic of the domain structures of SRSF1 and SRSF9. RRM, RNA recognition motif; RS, serine-arginine-rich region. (h, i) Localization of FLAG-SRSF9 WT or ΔRRM2 to HSATIII-scaffolded nSBs (h) and quantification of SRSF9-HSATIII colocalization using Pearson’s correlation coefficients (R) within nuclei (i). Nuclei were stained with DAPI. Scale bars, 10 μm. n = 30. Mann-Whitney U test was used. **** *P* < 0.0001. (k, l) Electrophoretic mobility shift assay (EMSA) of FITC-labeled GGAAU-repeat RNA with increasing concentrations of recombinant GST-SRSF9 RRM1 or RRM2 (k) and quantification of the ratio of shifted to free RNA (l). Data are shown as mean ± SD. n = 3. Šídák’s multiple-comparisons test was used. **** *P* < 0.0001.

**Extended Data Fig. 6.**
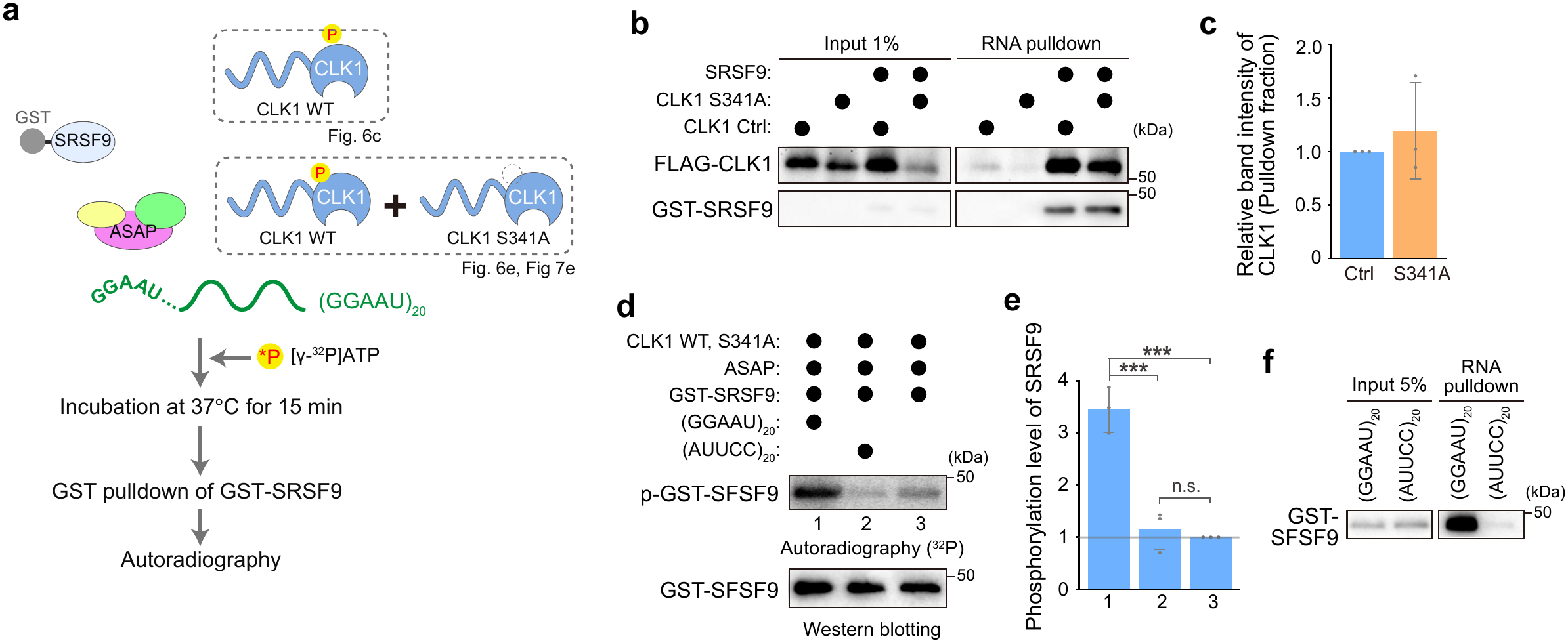
Biochemical characterization of nSB reaction-crucible activity. (a) Schematic of the in vitro SRSF9 phosphorylation assay using recombinant SRSF9, ASAP, and CLK1, together with GGAAU-repeat RNA. CLK1 was either WT phosphorylated Ser341 alone (d) or a mixture of WT phosphorylated at Ser341 and the unphosphorylated S341A mutant (f, and Fig. 7e). (b, c) In vitro RNA pulldown using recombinant SRSF9 and CLK1 (Ctrl and S341A) (b), and quantification of band intensity of CLK1 (c). Data are shown as mean ± SD. n = 3. Two-tailed t test was used. No statistically significant difference was observed. (d, e) In vitro phosphorylation of SRSF9 in the presence of GGAAU-repeat RNA or its antisense AUUCC repeat RNA, or in the absence of RNA (d). Quantification of SRSF9 phosphorylation (e). Šídák’s multiple-comparisons test was used. *** *P* < 0.001. Data are shown as mean ± SD. n = 3. (f) In vitro RNA pulldown of SRSF9 using GGAAU-repeat RNA and its antisense sequence, AUUCC, as baits.

**Extended Data Fig. 7.**
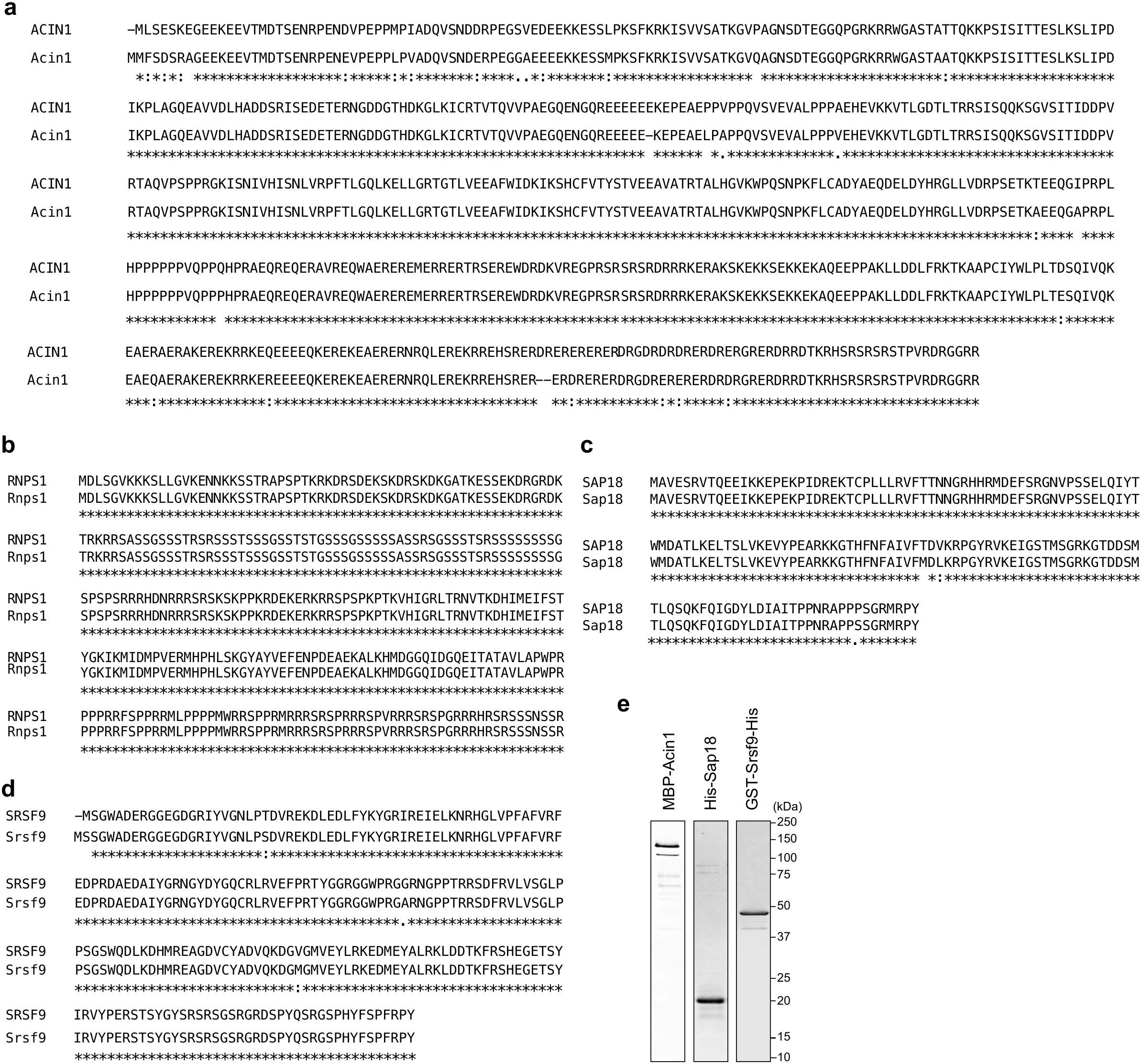
Evolutionary conservation of SRSF9 and ASAP components between humans and mice. (a-d) Amino acid sequence alignments of human and mouse ACIN1 (a), RNPS1 (b), SAP18 (c), and SRSF9 (d). (e) SDS-PAGE and CBB staining of recombinant mouse Acin1, Sap18, and Srsf9 used for in vitro assays.

**Supplementary Table 1. Hits identified by co-IP-MS screening.** Highlighted proteins indicate overlapping candidates. Proteins shown in red are components of the ASAP and PSAP complexes.

**Supplementary Table 2. The plasmids used in this study**

**Supplementary Table 3. The primers used in this paper**

**Supplementary Table 4. Antibodies used in this study**

**Supplementary Table 5. Oligonucleotides used in this study**

## References

1. Banani, S. F., Lee, H. O., Hyman, A. A. & Rosen, M. K. Biomolecular condensates: Organizers of cellular biochemistry. Nat Rev Mol Cell Biol. 18, 285–298 (2017).

2. Hirose, T., Ninomiya, K., Nakagawa, S. & Yamazaki, T. A guide to membraneless organelles and their various roles in gene regulation. Nat. Rev. Mol. Cell Biol. 24, 288–304 (2023).

3. Chujo, T. et al. Unusual semi-extractability as a hallmark of nuclear body-associated architectural noncoding RNA s . EMBO J. 36, 1447–1462 (2017).

4. Yamazaki, T. et al. Functional domains of NEAT1 architectural lncRNA induce paraspeckle assembly through phase separation. Mol. Cell 70, 1038–1053.e7 (2018).

5. Fujiwara, N., Ueno, T., Yamazaki, T. & Hirose, T. Unraveling architectural RNAs: Structural and functional blueprints of membraneless organelles and strategies for genome-scale identification. Biochim. Biophys. Acta Gen. Subj. 1869, (2025).

6. Hirose, T. et al. Architectural RNAs: blueprints for functional membraneless organelle assembly. Trends in Genetics https://doi.org/10.1016/j.tig.2025.05.007 (2025) doi:10.1016/j.tig.2025.05.007.

7. Shin, Y. & Brangwynne, C. P. Liquid phase condensation in cell physiology and disease. Science *(1979).* 357, (2017).

8. Mähl, P., Lutz, Y., Puvion, E. & Fuchs, J.-P. Rapid effect of heat shock on two heterogeneous nuclear ribonucleoprotein-associated antigens in HeLa cells. J. Cell Biol. 109, 1921–1935 (1989).

9. Jolly, C. et al. Stress-induced transcription of satellite III repeats. Journal of Cell Biology 164, 25–33 (2004).

10. Metz, A., Soret, J., Vourc’h, C., Tazi, J. & Jolly, C. A key role for stress-induced satellite III transcripts in the relocalization of splicing factors into nuclear stress granules. J. Cell Sci. 117, 4551–4558 (2004).

11. Rizzi, N. et al. Transcriptional activation of a constitutive heterochromatic domain of the human genome in response to heat shock. Mol. Biol. Cell 15, 543–551 (2004).

12. Ninomiya, K. et al. LncRNA -dependent nuclear stress bodies promote intron retention through SR protein phosphorylation . EMBO J. 39, (2020).

13. Denegri, M. et al. Human chromosomes 9, 12, and 15 contain the nucleation sites of stress-induced nuclear bodies. Mol. Biol. Cell 13, 2069–2079 (2002).

14. Jolly, C. & Lakhotia, S. C. Human sat III and Drosophila hsrω transcripts: A common paradigm for regulation of nuclear RNA processing in stressed cells. Nucleic Acids Res. 34, 5508–5514 (2006).

15. Jarmuz, M., Glotzbach, C. D., Bailey, K. A., Bandyopadhyay, R. & Shaffer, L. G. The evolution of satellite III DNA subfamilies among primates. Am. J. Hum. Genet. 80, 495–501 (2007).

16. Biamonti, G. & Vourc’h, C. Nuclear stress bodies. *Cold Spring Harb*. Perspect. Biol. 2, (2010).

17. Nurk, S. et al. The complete sequence of a human genome. Science *(1979).* 376, 44–3 (2022).

18. Ninomiya, K., Yamazaki, T. & Hirose, T. Satellite RNAs : emerging players in subnuclear architecture and gene regulation . EMBO J. 42, (2023).

19. Rhie, A. et al. The complete sequence of a human Y chromosome. Nature 621, 344–354 (2023).

20. Denegri, M. et al. Stress-induced nuclear bodies are sites of Accumulation of pre-mRNA processing factors. Mol. Biol. Cell 12, 3502–3514 (2001).

21. Ninomiya, K. & Hirose, T. Short tandem repeat-enriched architectural RNAs in nuclear bodies: Functions and associated diseases. Noncoding RNA 6, (2020).

22. Ninomiya, K. et al. m^6^A modification of HSATIII lncRNAs regulates temperature-dependent splicing . EMBO J. 40, (2021).

23. Chanseok Shin, Ying Feng & James L. Manley. Dephosphorylated SRp38 acts as a splicing repressor in response to heat shock. Nature 427, 548–553 (2004).

24. Ninomiya, K., Kataoka, N. & Hagiwara, M. Stress-responsive maturation of Clk1/4 pre-mRNAs promotes phosphorylation of SR splicing factor. Journal of Cell Biology 195, 27–40 (2011).

25. Shalgi, R., Hurt, J. A., Lindquist, S. & Burge, C. B. Widespread inhibition of posttranscriptional splicing shapes the cellular transcriptome following heat shock. Cell Rep. 7, 1362–1370 (2014).

26. Ueno, T. et al. Thermo-sensing mechanisms of splicing control by nuclear stress bodies. Mol. Cell 86, 3449–3465.e8 (2026).

27. Schwerk, C. et al. ASAP, a novel protein complex involved in RNA processing and apoptosis. Mol. Cell. Biol. 23, 2981–2990 (2003).

28. Ouyang, Y. et al. Discovery of 8-(6-Methoxypyridin-3-yl)-1-(4-(piperazin-1-yl)-3-(trifluoromethyl)phenyl)-1,5-dihydro-4H-[1,2,3]triazolo[4,5-c]quinolin-4-one (CQ211) as a highly potent and selective RIOK2 inhibitor. J. Med. Chem. 65, 7833–7842 (2022).

29. Murachelli, A. G., Ebert, J., Basquin, C., Le Hir, H. & Conti, E. The structure of the ASAP core complex reveals the existence of a Pinin-containing PSAP complex. Nat. Struct. Mol. Biol. 19, 378–386 (2012).

30. Mayeda, A. et al. Purification and characterization of human RNPS1: a general activator of pre-mRNA splicing. EMBO J. 18, 4560–4570 (1999).

31. Jens Lykke-Andersen, Mei-Di Shu & Joan A. Steitz. Communication of the position of exon-exon junctions to the mRNA surveillance machinery by the protein RNPS1. Science *(1979).* 293, 1836–1839 (2001).

32. Tange, T., Shibuya, T., Jurica, M. S. & Moore, M. J. Biochemical analysis of the EJC reveals two new factors and a stable tetrameric protein core. RNA 11, 1869– 1883 (2005).

33. Deka, B. & Singh, K. K. Multifaceted regulation of gene expression by the apoptosis- and splicing-associated protein complex and its components. Int. J. Biol. Sci. 13, 545–560 (2017).

34. Wang, Z., Ballut, L., Barbosa, I. & Le Hir, H. Exon Junction Complexes can have distinct functional flavours to regulate specific splicing events. Sci. Rep. 8, (2018).

35. Wang, F., Soprano, K. J. & Soprano, D. R. Role of acinus in regulating retinoic acid-responsive gene Pre-mRNA splicing. J. Cell. Physiol. 230, 791–801 (2015).

36. Setsuko S et al. Acinus is a caspase-3-activated protein required for apoptotic chromatin condensation. 401, 168–173 (1999).

37. Vohhodina, J. et al. The RNA processing factors THRAP3 and BCLAF1 promote the DNA damage response through selective mRNA splicing and nuclear export. Nucleic Acids Res. 45, 12816–12833 (2017).

38. Chiodi, I. et al. RNA recognition motif 2 directs the recruitment of SF2/ASF to nuclear stress bodies. Nucleic Acids Res. 32, 4127–4136 (2004).

39. Nishi, H., Hashimoto, K. & Panchenko, A. R. Phosphorylation in protein-protein binding: Effect on stability and function. Structure 19, 1807–1815 (2011).

40. Nishi, H., Fong, J. H., Chang, C., Teichmann, S. A. & Panchenko, A. R. Regulation of protein-protein binding by coupling between phosphorylation and intrinsic disorder: Analysis of human protein complexes. Mol. Biosyst. 9, 1620–1626 (2013).

41. Nishi, H., Shaytan, A. & Panchenko, A. R. Physicochemical mechanisms of protein regulation by phosphorylation. Front. Genet. 5, 270 (2014).

42. Cléry, A. et al. Isolated pseudo-RNA-recognition motifs of SR proteins can regulate splicing using a noncanonical mode of RNA recognition. Proc. Natl. Acad. Sci. U. S. A. 110, (2013).

43. Huang, H. et al. Tissue-selective restriction of RNA editing of CaV1.3 by splicing factor SRSF9. Nucleic Acids Res. 46, 7323–7338 (2018).

44. Hezroni, H. et al. Principles of long noncoding RNA evolution derived from direct comparison of transcriptomes in 17 species. Cell Rep. 11, 1110–1122 (2015).

45. Ulitsky, I. Evolution to the rescue: Using comparative genomics to understand long non-coding RNAs. Nat. Rev. Genet. 17, 601–614 (2016).

46. Mattick, J. S. et al. Long non-coding RNAs: definitions, functions, challenges and recommendations. Nat. Rev. Mol. Cell Biol. 24, 430–447 (2023).

47. Taniguchi, I. The SARS-CoV-2 ORF6 protein inhibits nuclear export of mRNA and spliceosomal U snRNA. PLoS One 19, (2024).

48. Rackham, O. & Brown, C. M. Visualization of RNA-protein interactions in living cells: FMRP and IMP1 interact on mRNAs. EMBO Journal 23, 3346–3355 (2004).

49. Takakuwa, H. et al. Shell protein composition specified by the lncRNA NEAT1 domains dictates the formation of paraspeckles as distinct membraneless organelles. Nat. Cell Biol. 25, 1664–1675 (2023).

