## Supplementary Table1, 2, 3, 4, 5 for "An architectural RNA builds a reaction crucible through molecular concentration and activated enzyme selection"

**Supplementary Table 1.** Hits identified by co-IP-MS screening. Highlighted proteins indicate overlapping candidates. Proteins shown in red are components of the ASAP and PSAP complexes.

| Protein | Log2 (CQ211/DMSO) | *p* value |
|---|---|---|
| CLASRP | -3.935 | 5.8.E-05 |
| **ACIN1** | **-3.167** | **2.8.E-05** |
| CFAP20 | -3.157 | 1.9.E-05 |
| MKLN1 | -3.122 | 2.0.E-02 |
| GPATCH8 | -3.028 | 3.4.E-06 |
| **RNPS1** | **-2.692** | **2.8.E-05** |
| NKTR | -2.438 | 3.8.E-06 |
| SRSF8 | -2.377 | 6.4.E-04 |
| PRMT1 | -1.993 | 6.8.E-09 |
| **SAP18** | **-1.945** | **2.0.E-04** |
| ERH | -1.844 | 3.8.E-04 |
| BCLAF1 | -1.843 | 1.8.E-03 |
| ZC3H18 | -1.829 | 8.3.E-05 |
| THRAP3 | -1.723 | 3.2.E-04 |
| GPALPP1 | -1.669 | 3.2.E-03 |
| PHRF1 | -1.652 | 3.2.E-03 |
| THOC5 | -1.634 | 2.6.E-02 |
| SUPT6H | -1.548 | 8.2.E-05 |
| DHX15 | -1.481 | 1.0.E-04 |
| NDUFA10 | -1.473 | 6.3.E-03 |
| DDX39A | -1.467 | 2.1.E-05 |
| RBBP6 | -1.437 | 1.1.E-05 |
| DDX39B | -1.405 | 1.3.E-05 |
| EIF4A2 | -1.338 | 6.8.E-04 |
| USP11 | -1.296 | 1.3.E-03 |
| PIR | -1.248 | 3.5.E-04 |
| ZRANB2 | -1.218 | 5.2.E-04 |
| SARNP | -1.202 | 1.6.E-04 |
| SYF2 | -1.199 | 1.0.E-03 |
| CBLL1 | -1.188 | 1.5.E-06 |
| LSM6 | -1.175 | 1.1.E-03 |
| ALDH3A2 | -1.170 | 1.1.E-04 |
| UBA2 | -1.162 | 3.4.E-03 |
| SRRT | -1.155 | 1.2.E-02 |
| HMGN5 | -1.106 | 2.2.E-02 |
| TIAL1 | 1.078 | 3.0.E-03 |
| REXO4 | -1.077 | 2.4.E-03 |
| LSM2 | -1.035 | 1.3.E-03 |
| UPF2 | -1.019 | 3.4.E-03 |
| **PNN** | **-1.009** | **3.3.E-03** |

| Protein | Log2 (ΔnSB-LRs/FL) | *p* value |
|---|---|---|
| AHCYL1 | -6.039 | 1.8E-06 |
| FMR1 | -5.607 | 1.0E-04 |
| AHCYL2 | -5.522 | 3.7E-05 |
| SARNP | -5.339 | 4.7E-06 |
| CASC3 | -5.066 | 4.3E-06 |
| WTAP | -4.523 | 6.5E-05 |
| LYZ | -4.483 | 1.1E-04 |
| PPIG | -4.324 | 1.4E-07 |
| **ACIN1** | **-4.176** | **2.8E-05** |
| DHX8 | -4.049 | 8.1E-06 |
| CPSF3 | -4.021 | 1.3E-08 |
| THOC5 | -3.976 | 1.0E-04 |
| CWC27 | -3.950 | 6.9E-04 |
| GPATCH8 | -3.935 | 3.6E-04 |
| SRRM1 | -3.904 | 2.3E-05 |
| CCDC9 | -3.903 | 8.7E-04 |
| **RNPS1** | **-3.809** | **1.5E-04** |
| SMNDC1 | -3.795 | 7.6E-05 |
| ZC3H14 | -3.769 | 2.2E-04 |
| CCDC12 | -3.756 | 3.8E-04 |
| THRAP3 | -3.745 | 5.3E-05 |
| BCLAF1 | -3.743 | 3.1E-04 |
| LUC7L3 | -3.718 | 3.2E-04 |
| WBP11 | -3.714 | 2.1E-03 |
| CLASRP | -3.694 | 2.3E-04 |
| SNRPC | -3.694 | 3.9E-04 |
| NCBP3 | -3.685 | 2.0E-04 |
| LSM2 | -3.678 | 2.2E-04 |
| LUC7L2 | -3.662 | 9.5E-04 |
| SYF2 | -3.612 | 2.3E-04 |
| CFAP20 | -3.608 | 6.7E-04 |
| ZC3H13 | -3.598 | 4.8E-05 |
| SNW1 | -3.597 | 2.0E-04 |
| **PNN** | **-3.585** | **1.1E-04** |
| THOC2 | -3.566 | 8.8E-06 |
| RBM22 | -3.549 | 1.1E-03 |
| FIP1L1 | -3.547 | 1.0E-03 |
| ERH | -3.545 | 1.7E-06 |
| ZCCHC3 | -3.533 | 7.1E-04 |
| MFAP1 | -3.518 | 2.0E-06 |
| DDX39A | -3.506 | 4.2E-04 |
| USP39 | -3.502 | 1.1E-04 |
| DDX39B | -3.491 | 1.5E-04 |
| NCBP1 | -3.489 | 8.1E-06 |
| PARP2 | -3.463 | 6.1E-05 |
| MAGOH | -3.425 | 1.2E-05 |
| AQR | -3.408 | 5.6E-04 |
| SF3A2 | -3.377 | 7.7E-05 |
| CRNKL1 | -3.329 | 8.0E-04 |
| JMJD6 | -3.327 | 4.9E-04 |
| THOC7 | -3.324 | 2.6E-03 |
| PRPF4B | -3.319 | 5.8E-06 |
| CPSF4 | -3.306 | 3.0E-04 |
| PRPF38A | -3.303 | 2.5E-04 |
| PAXBP1 | -3.302 | 1.2E-04 |
| RBM8A | -3.282 | 1.9E-04 |
| VIRMA | -3.271 | 6.0E-04 |
| RBM15 | -3.263 | 6.0E-04 |
| SRSF7 | -3.259 | 1.0E-04 |
| LSM4 | -3.257 | 1.1E-03 |
| EIF4A3 | -3.239 | 1.7E-04 |
| THOC1 | -3.229 | 2.7E-04 |
| THOC3 | -3.195 | 1.5E-03 |
| RBM15B | -3.183 | 1.6E-03 |
| CWC22 | -3.164 | 2.9E-03 |
| ZC3H11A | -3.141 | 1.5E-03 |
| RBM25 | -3.137 | 9.4E-06 |
| ALYREF | -3.127 | 1.0E-04 |
| CHTOP | -3.110 | 2.0E-04 |
| BUD31 | -3.107 | 3.3E-03 |
| SCAF1 | -3.090 | 1.4E-03 |
| POLDIP3 | -3.072 | 8.3E-04 |
| PPIL1 | -3.072 | 8.6E-04 |
| TRA2B | -3.068 | 4.3E-05 |
| THOC6 | -3.066 | 2.5E-05 |

| | | |
|---|---|---|
| CAAP1 | -3.060 | 3.0E-03 |
| MKRN2 | -3.049 | 1.8E-03 |
| U2AF2 | -3.044 | 5.1E-05 |
| **SAP18** | **-3.028** | **1.3E-04** |
| SRRT | -2.962 | 2.7E-05 |
| SNRPE | -2.948 | 1.0E-03 |
| XAB2 | -2.941 | 3.6E-03 |
| SF3A1 | -2.930 | 1.6E-04 |
| SNRPA | -2.927 | 2.1E-03 |
| PPIE | -2.924 | 4.5E-06 |
| SF3A3 | -2.920 | 1.3E-03 |
| LSM8 | -2.913 | 3.3E-03 |
| SNRNP40 | -2.909 | 1.3E-03 |
| FXR2 | -2.905 | 9.9E-06 |
| SRSF4 | -2.891 | 2.1E-05 |
| AFDN | -2.880 | 2.4E-02 |
| SNRPF | -2.868 | 1.9E-03 |
| NCBP2 | -2.865 | 1.1E-04 |
| ZCCHC8 | -2.865 | 9.3E-06 |
| SNRPG | -2.817 | 2.0E-03 |
| PABPN1 | -2.813 | 1.6E-03 |
| SLTM | -2.789 | 6.5E-04 |
| UPF3B | -2.784 | 1.1E-02 |
| BRPF3 | -2.784 | 1.2E-03 |
| NCOA5 | -2.774 | 4.3E-03 |
| KRT73 | -2.750 | 2.9E-02 |
| SRSF10 | -2.733 | 4.3E-04 |
| SRSF6 | -2.722 | 1.9E-03 |
| CPSF1 | -2.717 | 1.6E-03 |
| SNRPA1 | -2.710 | 1.7E-03 |
| TRA2A | -2.708 | 5.5E-06 |
| DHX38 | -2.706 | 3.0E-04 |
| SMU1 | -2.692 | 1.2E-03 |
| EIF3B | -2.685 | 2.3E-03 |
| CWC15 | -2.678 | 2.6E-03 |
| SNRPD1 | -2.670 | 1.3E-03 |
| LSM7 | -2.667 | 1.2E-03 |
| SPTY2D1 | -2.636 | 1.9E-04 |
| SNRPD2 | -2.626 | 9.1E-04 |
| WBP4 | -2.625 | 1.7E-02 |
| SNRPD3 | -2.613 | 2.1E-03 |
| FAM120C | -2.613 | 2.5E-03 |
| PRPF8 | -2.593 | 1.2E-03 |
| FXR1 | -2.592 | 3.8E-05 |
| MTERF3 | -2.581 | 3.8E-02 |
| CPSF2 | -2.573 | 4.1E-03 |
| TDRD3 | -2.571 | 4.1E-03 |
| PRPF6 | -2.567 | 4.4E-03 |
| SAFB2 | -2.567 | 4.7E-04 |
| SETD2 | -2.527 | 1.0E-05 |
| SRSF8 | -2.504 | 8.0E-03 |
| SNRNP200 | -2.493 | 1.2E-03 |
| CSTF3 | -2.489 | 2.2E-02 |
| IK | -2.475 | 5.2E-03 |
| DDX41 | -2.471 | 1.6E-03 |
| PRMT1 | -2.447 | 1.3E-03 |
| SCAF11 | -2.440 | 6.7E-04 |
| RBMX | -2.426 | 1.8E-04 |
| ERP44 | -2.424 | 4.9E-02 |
| HNRNPK | -2.411 | 2.4E-03 |
| PRRC2B | -2.408 | 9.7E-04 |
| TOP3B | -2.401 | 9.2E-03 |
| SNRPB | -2.355 | 3.5E-03 |
| WDR33 | -2.355 | 2.0E-02 |
| PABPC4 | -2.327 | 1.2E-04 |
| HNRNPC | -2.312 | 8.6E-04 |
| MRPL10 | -2.296 | 5.9E-03 |
| SNRNP70 | -2.290 | 7.5E-03 |
| PRPF31 | -2.288 | 8.6E-05 |
| SRRM2 | -2.264 | 6.7E-04 |
| U2AF1 | -2.257 | 5.6E-04 |
| ELAVL1 | -2.225 | 3.3E-03 |
| PPRC1 | -2.223 | 1.8E-03 |
| PABPC1 | -2.223 | 3.4E-04 |
| DCD | -2.211 | 2.6E-02 |
| PPHLN1 | -2.195 | 2.4E-02 |
| EIF3CL | -2.193 | 1.2E-05 |
| CDC40 | -2.185 | 3.2E-03 |
| TFIP11 | -2.180 | 1.0E-03 |

| | | |
|---|---|---|
| MRPS11 | -2.180 | 2.4E-03 |
| SRSF1 | -2.171 | 3.9E-03 |
| MTREX | -2.150 | 1.0E-05 |
| SNRPB2 | -2.148 | 6.7E-03 |
| RALY | -2.145 | 3.1E-03 |
| PPWD1 | -2.142 | 3.1E-03 |
| ISY1 | -2.139 | 8.7E-03 |
| CSNK2A2 | -2.133 | 3.6E-03 |
| SF3B5 | -2.130 | 1.6E-03 |
| EIF3L | -2.129 | 9.6E-03 |
| SRSF2 | -2.122 | 2.2E-03 |
| CELF1 | -2.121 | 9.4E-03 |
| HNRNPCL1 | -2.113 | 3.1E-04 |
| SF3B1 | -2.104 | 2.5E-03 |
| YTHDC1 | -2.101 | 2.2E-02 |
| IMP3 | -2.092 | 1.6E-03 |
| IGF2BP1 | -2.035 | 7.7E-04 |
| NKAP | -2.016 | 8.5E-05 |
| MRPS33 | -2.004 | 9.8E-04 |
| MRPL48 | -1.995 | 7.6E-03 |
| HNRNPL | -1.992 | 5.0E-03 |
| NKTR | -1.988 | 4.8E-03 |
| POLR2B | -1.965 | 1.1E-02 |
| SF3B6 | -1.935 | 4.5E-03 |
| EIF3M | -1.935 | 1.9E-02 |
| MRPS14 | -1.933 | 4.9E-04 |
| CCAR2 | -1.924 | 1.1E-02 |
| EIF3I | -1.924 | 1.7E-02 |
| ZRANB2 | -1.920 | 4.2E-02 |
| MRPS31 | -1.918 | 2.0E-04 |
| CTNNBL1 | -1.913 | 1.8E-02 |
| SFMBT1 | -1.908 | 2.7E-03 |
| FYTTD1 | -1.906 | 4.0E-03 |
| LARP1 | -1.904 | 1.0E-03 |
| ATXN2 | -1.888 | 6.0E-04 |
| MRPS10 | -1.879 | 1.6E-03 |
| TIAL1 | -1.851 | 2.0E-03 |
| FLG2 | -1.848 | 4.7E-02 |
| EIF3E | -1.842 | 2.9E-02 |
| PRPF40A | -1.837 | 4.1E-04 |
| PEG10 | -1.836 | 4.5E-02 |
| UPF1 | -1.834 | 3.1E-03 |
| MRPS2 | -1.834 | 3.2E-04 |
| ANK2 | -1.820 | 2.1E-02 |
| PTCD3 | -1.808 | 1.9E-03 |
| CD2BP2 | -1.808 | 2.1E-02 |
| MRPS18C | -1.807 | 4.8E-03 |
| MRPS35 | -1.797 | 8.1E-04 |
| SF3B3 | -1.792 | 9.7E-03 |
| PPIL3 | -1.789 | 5.4E-03 |
| UPF2 | -1.782 | 1.8E-03 |
| NFX1 | -1.778 | 2.6E-02 |
| EIF3F | -1.772 | 2.4E-05 |
| MRPS15 | -1.772 | 1.4E-03 |
| ZC3H18 | -1.765 | 1.0E-02 |
| IGF2BP3 | -1.741 | 3.6E-03 |
| SNIP1 | -1.740 | 2.2E-04 |
| MRPS21 | -1.738 | 1.6E-04 |
| MRPS27 | -1.732 | 2.3E-03 |
| PARP1 | -1.731 | 2.4E-02 |
| RBBP6 | -1.727 | 3.8E-03 |
| SRSF5 | -1.716 | 4.1E-03 |
| DHX29 | -1.701 | 4.5E-03 |
| PUF60 | -1.688 | 3.7E-03 |
| RBM17 | -1.682 | 2.6E-03 |
| ZNF326 | -1.679 | 6.6E-03 |
| FAM120A | -1.677 | 6.8E-03 |
| MRPS24 | -1.675 | 4.9E-03 |
| CSNK2A1 | -1.668 | 1.1E-03 |
| MRPS16 | -1.666 | 4.4E-05 |
| MRPS5 | -1.656 | 2.4E-04 |
| MRPS18B | -1.654 | 2.5E-03 |
| SON | -1.643 | 1.2E-04 |
| MRPS28 | -1.639 | 1.9E-03 |
| EIF3A | -1.630 | 5.3E-03 |
| DAP3 | -1.622 | 4.1E-05 |
| MRPS7 | -1.619 | 7.9E-04 |
| MRPS23 | -1.611 | 2.6E-05 |
| ATXN2L | -1.608 | 2.2E-03 |

| | | |
|---|---|---|
| MRPS6 | -1.583 | 5.9E-04 |
| PHF5A | -1.554 | 4.5E-02 |
| TRIP4 | -1.549 | 5.7E-07 |
| HNRNPA3 | -1.546 | 1.1E-02 |
| EIF3H | -1.538 | 2.2E-02 |
| GCN1 | -1.535 | 4.2E-03 |
| C1orf35 | -1.530 | 1.5E-04 |
| MRPS34 | -1.518 | 1.7E-06 |
| PPIH | -1.509 | 3.7E-03 |
| LARP4 | -1.508 | 5.7E-03 |
| KPNA4 | -1.500 | 5.9E-03 |
| PRPF4 | -1.491 | 7.7E-03 |
| PURA | -1.489 | 3.7E-04 |
| PRPF3 | -1.473 | 1.8E-02 |
| DDX3X | -1.469 | 1.2E-02 |
| MRPS9 | -1.468 | 4.2E-04 |
| MRPS26 | -1.466 | 1.0E-04 |
| SF3B2 | -1.456 | 5.1E-03 |
| EFTUD2 | -1.449 | 4.8E-04 |
| MBNL1 | -1.440 | 8.7E-03 |
| HNRNPR | -1.437 | 7.6E-03 |
| DHX15 | -1.431 | 4.9E-02 |
| HNRNPUL2 | -1.431 | 2.1E-02 |
| MIS18A | -1.430 | 7.2E-05 |
| ILF3 | -1.427 | 4.8E-03 |
| SAFB | -1.401 | 8.6E-05 |
| FUBP3 | -1.393 | 1.5E-02 |
| ILF2 | -1.376 | 6.2E-03 |
| AKAP8 | -1.376 | 4.1E-02 |
| MRPS22 | -1.361 | 5.3E-04 |
| HNRNPAB | -1.360 | 1.8E-02 |
| SF3B4 | -1.344 | 7.6E-03 |
| SRSF11 | -1.343 | 2.3E-05 |
| EIF4G1 | -1.340 | 3.0E-02 |
| COQ6 | -1.314 | 7.7E-04 |
| FAM98A | -1.311 | 9.2E-05 |
| SRSF9 | -1.305 | 5.4E-03 |
| EIF4E | -1.299 | 7.0E-03 |
| PTBP1 | -1.297 | 1.5E-02 |
| AKAP17A | -1.293 | 2.2E-02 |
| NAF1 | -1.286 | 3.7E-02 |
| G3BP1 | -1.279 | 6.6E-04 |
| DDX17 | -1.278 | 9.6E-03 |
| EIF3D | -1.278 | 2.3E-02 |
| GP2 | -1.255 | 9.1E-03 |
| C14orf166 | -1.250 | 7.0E-04 |
| MRPS17 | -1.241 | 9.1E-04 |
| CHERP | -1.240 | 8.0E-03 |
| SRSF3 | -1.236 | 1.6E-06 |
| CAPRIN1 | -1.222 | 1.8E-04 |
| PRRC2A | -1.214 | 1.1E-02 |
| IGF2BP2 | -1.210 | 3.3E-03 |
| SART1 | -1.196 | 1.4E-02 |
| DHX36 | -1.181 | 1.3E-02 |
| LSM12 | -1.166 | 6.3E-03 |
| YBX1 | -1.156 | 7.0E-03 |
| PHAX | -1.153 | 2.5E-02 |
| MOV10 | -1.151 | 6.3E-03 |
| MRPS25 | -1.143 | 8.1E-05 |
| HNRNPH1 | -1.139 | 2.7E-02 |
| YBX3 | -1.137 | 1.5E-03 |
| PCBP2 | -1.119 | 3.4E-02 |
| NUFIP2 | -1.107 | 1.9E-04 |
| CSNK2B | -1.095 | 6.1E-08 |
| PNKD | -1.092 | 3.2E-03 |
| G3BP2 | -1.086 | 2.2E-03 |
| RTCB | -1.075 | 7.6E-03 |
| PTBP3 | -1.075 | 9.5E-04 |
| GKAP1 | -1.069 | 1.5E-04 |
| DDX5 | -1.069 | 1.9E-02 |
| SORCS2 | -1.048 | 4.7E-03 |
| CT45A1 | -1.048 | 4.1E-04 |
| DDX1 | -1.029 | 8.7E-05 |
| U2SURP | -1.026 | 3.6E-04 |
| MGAT3 | -1.019 | 1.5E-03 |
| CEP41 | -1.006 | 8.3E-04 |

**Supplementary Table 2.** The plasmids used in this study.

| Plasmids | Sorce | Identifier |
|---|---|---|
| pcDNA5.FRT.TO.i.3xFLAG-CLK1.K191R | Ninomiya et al., *EMBO J* , 2020 | N/A |
| pcDNA5.FRT.TO.i.3xFLAG-CLK1.K191R, S341A | Ueno et al., *Mol Cell* , 2026 | N/A |
| pcDNA5.FRT.TO.i.3xFLAG-CLK1.S341A | Ueno et al., *Mol Cell* , 2026 | N/A |
| pcDNA5.FRT.TO.i.3xFLAG-CLK1.WT | Ninomiya et al., *EMBO J* , 2020 | N/A |
| pcDNA5.FRT.TO.i.3xHA-ACIN1-L.siRNA#2_resisitant | This paper | N/A |
| pcDNA5.FRT.TO.i.3xHA-ACIN1-S | This paper | N/A |
| pcDNA5.FRT.TO.i.3xHA-ACIN1-S ΔRRM | This paper | N/A |
| pcDNA5.FRT.TO.i.3xHA-ACIN1-S ΔRS | This paper | N/A |
| pcDNA5.FRT.TO.i.3xHA-ACIN1-S ΔRSB | This paper | N/A |
| pcDNA5.FRT.TO.i.3xHA-ACIN1-S ΔRSB.siRNA#2_resisitant | This paper | N/A |
| pcDNA5.FRT.TO.i.3xHA-ACIN1-S.siRNA#2_resisitant | This paper | N/A |
| pcDNA5.FRT.TO.i.3xHA-PNN | This paper | N/A |
| pcDNA5.FRT.TO.i.3xHA-RNPS1 | This paper | N/A |
| pcDNA5.FRT.TO.i.3xHA-RNPS1 ΔC | This paper | N/A |
| pcDNA5.FRT.TO.i.3xHA-RNPS1 ΔN | This paper | N/A |
| pcDNA5.FRT.TO.i.3xHA-RNPS1 ΔRRM | This paper | N/A |
| pcDNA5.FRT.TO.i.3xHA-RNPS1 ΔRRM.siRNA#2_resisitant | This paper | N/A |
| pcDNA5.FRT.TO.i.3xHA-RNPS1.siRNA#2_resisitant | This paper | N/A |
| pcDNA5.FRT.TO.i.3xHA-SAP18 | This paper | N/A |
| pcDNA5.FRT.TO.i.EGFP-CLK1.K191R | Ueno et al., *Mol Cell* , 2026 | N/A |
| pcDNA5.FRT.TO.i.EGFP-CLK1.K191R, S341A | This paper | N/A |
| pcDNA5.FRT.TO.i.MBP-ACIN1-S | This paper | N/A |
| pcDNA5.FRT.TO.i.MBP-Acin1-S | This paper | N/A |
| pcDNA5.FRT.TO.i.MBP-ACIN1-S ΔRSB | This paper | N/A |
| pcDNA5.FRT.TO.i.MBP-RNPS1 | This paper | N/A |
| pcDNA5.FRT.TO.i.MBP-RNPS1 ΔRRM | This paper | N/A |
| pcDNA3.FLAG-SRSF9 WT | Ninomiya et al., *EMBO J* , 2020 | N/A |
| pcDNA3.FLAG-SRSF9 ΔRRM2 | This paper | N/A |
| pET28a-6xHis-SAP18 | This paper | N/A |
| pET28a-6xHis-Sap18 | This paper | N/A |
| pET28a-6xHis-SRSF1 | This paper | N/A |
| pGEX.6p-1.GST-Empty-6xHis | Taniguchi, *Plos One* , 2024 | N/A |
| pGEX.6p-1.GST-RIOK2-6xHis | This paper | N/A |
| pGEX.6p-1.GST-SRSF9 RRM1-6xHis | This paper | N/A |
| pGEX.6p-1.GST-SRSF9 RRM2-6xHis | This paper | N/A |
| pGEX.6p-1.GST-SRSF9 ΔRRM1-6xHis | This paper | N/A |
| pGEX.6p-1.GST-SRSF9 ΔRRM2-6xHis | This paper | N/A |
| pGEX.6p-1.GST-SRSF9 ΔRS-6xHis | This paper | N/A |
| pGEX.6p-1.GST-SRSF9-6xHis | This paper | N/A |
| pGEX.6p-1.GST-Srsf9-6xHis | This paper | N/A |
| pGEX.6p-1.MBP-FLAG-CLK1.K191R-6xHis | This paper | N/A |
| pGEX.6p-1.MBP-FLAG-Empty-6xHis | Taniguchi, *Plos One* , 2024 | N/A |
| pGEX.6p-1.MBP-FLAG-RNPS1-6xHis | This paper | N/A |
| pLVSIN-FLAG-MCP | This paper | N/A |
| pLVSIN-FLAG-MCP-ACIN1-L | This paper | N/A |
| pLVSIN-FLAG-MCP-RNPS1 | This paper | N/A |
| pLVSIN-FLAG-MCP-SAP18 | This paper | N/A |

**Supplementary Table 3**. The primers used in this paper.

| Primers | Forward | Reverse |
|---|---|---|
| **Primers for plasmid construction (5’→3’)** | | |
| ACIN1-L (BamHI, NotI) | AAAAGGATCCATGTGGAGACGGAAACATCCGAGGAC | GGGGGGCGGCCGCCTAGCGGCGCCCACCCCGGTCCCGCAC |
| ACIN1-S (BamHI, NotI) | AAAAGGATCCATGTTATCAGAAAGCAAAGAAGGTG | AAAAGGATCCATGTTATCAGAAAGCAAAGAAGGTG |
| Acin1-S (BamHI, XhoI) | AAAAAGGATCCATGATGTTCTCAGACAGCAGAGCAGGTG | GGGGGCTCGAGCTAGCGGCGCCCACCCCGGTCCCGTACAG |
| CLK1 (BamHI, XhoI) | TCAGACGGATCCATGAGACACTCAAAGAGAACTTAC | AATCTGCTCGAGAGAGCTGTCCAATTACAG |
| CLK1 Δ1 mutation (BamHI, XhoI) | AAAAGGATCCATGAAAAGAAGGAAGAGATCAC | AATCTGCTCGAGAGAGCTGTCCAATTACAG |
| PNN (XhoI, ApaI) | AAAAACTCGAGGATGGCGGTCGCCGTGAGAACTTTGCAG | CCCCCGGGCCCTTAACGCCTTTTGTCTTTCCTGTCTG |
| RIOK2 (BamHI, XhoI) | AAAAGGATCCATGGGGAAAGTGAATGTGGCCAAGTTGC | GGGGGCTCGAGTTATTCTCCCCAAAAGCTGGCTG |
| RNPS1 (BamHI, XhoI) | AAAAGGATCCATGGATTTATCAGGAGTGAAAAAGAAGAGC | GGGGGCTCGAGTTATCGGGAGGAGTTGGAGCTGGAGC |
| SAP18 (BamHI, XhoI) | AAAAGGATCCATGGCGGTGGAGTCGCGCGTTACC | GGGGGCTCGAGTTAATATGGTCTCATGCGCCCTGAAG |
| Sap18 (BamHI, XhoI) | AAAAAGGATCCATGGCGGTGGAGTCGCGCGTTACCCAG | GGGGGCTCGAGTCAGTAGGGTCTCATCCTCCCTGACG |
| SRSF1 (BamHI, XhoI) | AAAAGGATCCATGTCGGGAGGTGGTGTGATTCG | GGGGGGCGGCCGCTTATGTACGAGAGCGAGATCTGC |
| SRSF1 (BamHI, XhoI) | AAAAGGATCCATGTCGGGAGGTGGTGTGATTCG | GGGGGGCGGCCGCTTATGTACGAGAGCGAGATCTGC |
| SRSF9 (BamHI, XhoI) | AAAAGGATCCATGTCGGGCTGGGCGGACGAG | GGGGGCTCGAGGTAGGGCCTGAAAGGAGAGAAGTAGTG |
| Srsf9 (In-fusion to pGEX.6p-1 vector) | TTCCAGGGGCCCCTGGGATCCATGTCGTCGGGCTGG | CTCAGTGGTGGTGGTGGTGGTGGTAGGGCCTGAAAGGAGAGAAG |
| SRSF9.RRM1 (BamHI, XhoI) | AAAAAGGATCCGGGCGCATCTACGTGGGGAACC | GGGGGCTCGAGAGTCCTGGGGAACTCCACACG |
| SRSF9.RRM2 (BamHI, XhoI) | AAAAGGATCCTTCCGAGTTCTTGTTTCAGGAC | GGGGCTCGAGGCTTCTCTCAGGATAAACTCGGATG |
| **Primers for PCR-based mutagenesis (5’→3’)** | | |
| ACIN1 siRNA.#2_resistant | TGACCTATAGAACAGTAGAGGAAGCTGTTGCCAC | CGAAACAGTGGCTTTTGATCTTGTCAATCCAGAAGG |
| ACIN1 ΔRRM | TCCGAACTAAGACAGAGGAGCAGGGAATACCACG | TCTGTCTTAGTTCGGACTGGGTCATCAATGGTAATGG |
| ACIN1 ΔRS-1 | AAGGGCCCAAGGAACGTGCGAAGTCTAAAGAAAAGAAGAG | ACGTTCCTTGGGCCCTTCTCGAACTTTGTCC |
| ACIN1 ΔRS-2 | AAGCGCCACGTGCGGGACCGGGGTGGGCGCCGCTAG | TCCCGCACGTGGCGCTTGGTGTCCCTGCGATCC |
| ACIN1 ΔRSB | AGAAAGCCCGGCGAAAGGAGCAAGAAGAAGAAGAGC | TTTCGCCGGGCTTTCTCTTTCTTCTCACTCTTC |
| CLK1 Δ2 mutation | GGAGGAGCAGCAGCAGTCATAATGAGAAAGATTATCATAGTC | GACTATGATAATCTTTCTCATTATGACTGCTGCTGCTCCTCC |
| CLK1 Δ3 mutation | CCATTATTTGGAAAGCAGGTCTATAAGTAGCAAGTCTTCTGG | CCAGAAGACTTGCTACTTATAGACCTGCTTTCCAAATAATGG |
| CLK1 Δ4 mutation | CCATGAAAGCCGGTATCAGAACCATCACCGAAGGAAAAGAACC | GGTTCTTTTCCTTCGGTGATGGTTCTGATACCGGCTTTCATGG |
| CLK1 Δ5 mutation | GAAATTGTTGATACTTTAGGTGAAGGAGCTTTTGG | ACTCTTCCCATGTGAACGACGATGTGAAGTACTG |
| CLK1 K191R mutant | GACATGTAGCAGTAAGAATAGTTAAAAATGTG | CACATTTTTAACTATTCTTACTGCTACATGTC |
| CLK1 S341A mutation | TGGTAGCCACAAGACATTATAGAGCACCTG | CTTGTGGCTACCAATGTACTGTGATGTTCGTC |
| RNPS1 siRNA.#2_resistant | AAGGCGGAGTCCTAGCCCAAAACCAACAAAGGTGCACATTGG | CCAATGTGCACCTTTGTTGGTTTTGGGCTAGGACTCCGCCTT |
| RNPS1 ΔC | GTGCTGGCCTAACTCGAGTCTAGAGGGCCCGTTTAAAC | ACTCGAGTTAGGCCAGCACGGCGGTGGCAGTGATCTC |
| RNPS1 ΔN | AGCTCGGATCCACCAAAGTGCACATTGGGAGACTCAC | TGCACTTTGGTGGATCCGAGCTCGGTACCAAGCTTAGC |
| RNPS1 ΔRRM | TCCTAAGCCCCCCTGGCCTAGGCCACCCCCCAGGAG | TAGGCCAGGGGGGCTTAGGAGATGGGCTCCGCCTTTTCC |
| RNPS1 ΔRRM siRNA.#2_resistant | CAAAACCACCCTGGCCTAGGCCACCCCCCAGG | GGCTAGGACTCCGCCTTTTCCTCTCCTTTTCATCTC |
| SRSF9 ΔRRM1 | AGGGCGACTATGGAGGTCGGGGTGGGTGGCCCCGTGG | ACCTCCATAGTCGCCCTCGCCGCCGCGCTCGTCC |
| SRSF9 ΔRRM2 | AAGATCTGATTTCTACATCCGAGTTTATCCTGAGAG | AAACTCGGATGTAGAAATCAGATCTTCTTGTAGGAGG |
| SRSF9 ΔRS | AGCACCAGCTGACTCGAGCATGCATCTAGAGGGCCCTATTC | TGCTCGAGTCAGCTGGTGCTTCTCTCAGGATAAACTCG |
| **Primers for qRT-PCR (5’→3’)** | | |
| ABHD5 intron 5 detained | ATGGTGCCCTACGTCTATATCAC | ACACCACAGAATCTTACCTCCAG |
| ABHD5 intron 5 spliced | ATGGTGCCCTACGTCTATATCAC | ATGGAGAAGGACAAGTGGAGTC |
| AP4B1 intron 3 detained | CCTGGCCATCAATACGCTGTGC | CATCAGCCATTTAGGACCAGACAGA |
| AP4B1 intron 3 spliced | CCTGGCCATCAATACGCTGTGC | AGCCTTATCCCGCAGACCATTG |
| CLK1 intron 3 detained | TTCACATCGTCGTTCACATGG | ATCCTGCAGGTTAAAACTGAGATG |
| CLK1 intron 3 spliced | TTCACATCGTCGTTCACATGG | ATCTTGCACTTAGTACGTCTC |
| DAB2 intron 6 detained | TGTTGACCAGATGGATTTGTTTGG | ATGCTTAAGATTTCTGGTTATTGCC |
| DAB2 intron 6 spliced | TGTTGACCAGATGGATTTGTTTGG | TGTCGATTTCAGAGTTTAGATCCAC |
| FAM214A intron 6 retaining□ | CTGTTTGTCTACATGTGAGCAACC | TTGGGCCTTTATAACCAGATCTCC |
| FAM214A intron 6 spliced□ | CTGTTTGTCTACATGTGAGCAACC | AGTTTGAGGATGAAAGGCTCTGG |
| MORC1 intron 1 detained | AGCAGTCTGAATCGAGCTCAGC | TTCACTCCAGTGGGATACATCTC |
| MORC1 intron 1 spliced | AGCAGTCTGAATCGAGCTCAGC | ACCAAACAAGAATTCGTGAGTGG |
| SDHA intron 11 detained | TGCTGATGGAAGCATAAGAACATCG | AACTGCACCTGCATCCCCCTAG |
| SDHA intron 11 spliced | TGCTGATGGAAGCATAAGAACATCG | TCTTCAGGTGCTTTAGGTCTCC |
| TAF1D intron 2 retaining | ATGTGACATCTGATGCTGTGG | TCACTTGCGTGAACACTTTCAGG |
| TAF1D intron 2 spliced | ATGTGACATCTGATGCTGTGG | ACTTGGAAGGTCAGTGAAATCTAG |
| TSR1 intron 5 detained | AAGGACCCAGACATGGCAATGG | ACTATTGCCTTACTTCAGACTAGC |
| TSR1 intron 5 spliced | AAGGACCCAGACATGGCAATGG | TCAGTGGGCCAGGTTTGCTCTC |
| **Templates for in vitro transcription (5’→3’)** | | |
| T7 promoter | CTAATACGACTCACTATAGGGAGA | |
| T7 promoter-GGAAT | ATTCCATTCCATTCCATTCCATTCCATTCCATTCCATTCCATTCCATTCCATTCCAT TCCATTCCATTCCATTCCATTCCATTCCATTCCATTCCATTCCTCTCCCTATAGTG AGTCGTATTAG | |
| T7 promoter-GGAAT AS | GGAATGGAATGGAATGGAATGGAATGGAATGGAATGGAATGGAATGGAATGGAA TGGAATGGAATGGAATGGAATGGAATGGAATGGAATGGAATGGAATTCTCCCTA TAGTGAGTCGTATTAG | |

**Supplementary Table 4.** Antibodies used in this study.

| Antibodies | Sorce | Identifier |
|---|---|---|
| Mouse monoclonal anti-FLAG (clone M2) | Sigma-Aldrich | Cat# F3165; RRID: AB_259529 |
| Mouse monoclonal anti-GAPDH (clone 6C5) | Santa Cruz Biotechnology | Cat# sc-32233; RRID: AB_627679 |
| Mouse monoclonal anti-GST-tag | Proteintech | Cat# 66001-2-Ig; RRID: AB_2881488 |
| Mouse monoclonal anti-HA.11 (clone 16B12) | BioLegend | Cat# 901515; RRID: AB_2565334 |
| Mouse monoclonal anti-His-tag | MBL | Cat# D291-3; RRID: AB_10597733 |
| Mouse monoclonal anti-MBP (R29.6) | santa | Cat# sc-13564; RRID: AB_675707 |
| Mouse monoclonal anti-SC35 | Sigma-Aldrich (Merck) | Cat# S4045; RRID: AB_477511 |
| Mouse monoclonal anti-SR (1H4) | Santa Cruz Biotechnology | Cat# sc-13509; RRID: AB_2185202 |
| Rabbit monoclonal anti-HA-tag (clone C29F4) | Cell signaling technology | Cat# 3724; RRID: AB_1549585 |
| Rabbit polyclonal anti-Acinus | Cell Signaling Technology | Cat# 4934; RRID: AB_2257955 |
| Rabbit polyclonal anti-alpha Tubulin | Abcam | Cat# ab15246; RRID: AB_301787 |
| Rabbit polyclonal anti-BCLAF1 | Atlas antibodies | Cat# HPA027770; RRID: AB_1845484 |
| Rabbit polyclonal anti-GAPDH | Abcam | Cat# ab37168; RRID: AB_732652 |
| Rabbit polyclonal anti-GFP | MBL | Cat# 598; RRID: AB_591819 |
| Rabbit polyclonal anti-Pinin | Proteintech | Cat# 18266-1-AP; RRID: AB_10642138 |
| Rabbit polyclonal anti-RNPS1 | GeneTex | Cat# GTX129789; RRID: AB_2886089 |
| Rabbit polyclonal anti-SAP18 | Proteintech | Cat# 13841-1-AP; RRID: AB_2301353 |
| Rabbit polyclonal anti-SON/DRP5 | Abcam | Cat# ab121759; RRID: AB_11132447 |
| Rabbit polyclonal anti-SRSF9 | MBL | Cat# RN081PW; RRID: AB_11160952 |
| Rabbit polyclonal anti-THRAP3 | Bethyl Laboratories | Cat# A300-956A; RRID: AB_805852 |
| Goat anti-Mouse IgG (H+L) Superclonal™ Secondary Antibody (Alexa Fluor™ 488) | Thermo Fisher Scientific | Cat# A28175; RRID: AB_2536161 |
| Goat anti-Rabbit IgG (H+L) Highly Cross-Adsorbed Secondary Antibody ( Alexa Fluor™ 488) | Thermo Fisher Scientific | Cat# A-11034; RRID: AB_2576217 |
| Goat anti-Mouse IgG (H+L) Highly Cross-Adsorbed Secondary Antibody ( Alexa Fluor™ 568) | Thermo Fisher Scientific | Cat# A-11031; RRID: AB_144696 |
| Goat anti-Rabbit IgG (H+L) Highly Cross-Adsorbed Secondary Antibody (Alexa Fluor™ 568) | Thermo Fisher Scientific | Cat# A-11036; RRID: AB_10563566 |
| Donkey polyclonal anti-Rabbit IgG (HRP-Linked) | Cytiva | Cat# NA934; RRID: AB_772206 |
| Sheep polyclonal anti-Mouse IgG (HRP-Linked) | Cytiva | Cat# NA931; RRID: AB_772210 |

**Supplementary Table 5**. Oligonucleotides used in this study.

| siRNAs | Sorce | Identifier |
|---|---|---|
| Control Stealth siRNA (Luciferase Stealth siRNA) (5′-CAAAUGCAGUUACUGGUGUUGAUUA-3′) | Ueno et al, *Mol Cell* , 2026 | N/A |
| ACIN1 Stealth siRNA | Thermo Fisher Scientific | HSS117988 |
| ACIN1 Stealth siRNA | Thermo Fisher Scientific | HSS117990 |
| PNN Stealth siRNA | Thermo Fisher Scientific | HSS108190 |
| PNN Stealth siRNA | Thermo Fisher Scientific | HSS108191 |
| RNPS1 Stealth siRNA | Thermo Fisher Scientific | HSS152719 |
| RNPS1 Stealth siRNA | Thermo Fisher Scientific | HSS152720 |
| RIOK2 Stealth siRNA | Thermo Fisher Scientific | HSS183267 |
| Silencer™ Select Negative Control No. 1 siRNA | Thermo Fisher Scientific | 4390843 |
| THRAP3 Silencer siRNA | Thermo Fisher Scientific | s19360 |
| BCLAF1 Silencer siRNA | Thermo Fisher Scientific | s18875 |
| **ASOs** | **Sorce** | **Identifier** |
| HSATIII ASO for ChIRP (5'-ATTCCATTCCATTCCATTCC-3', 3' biotin-labeled) | Ninomiya et al., *EMBO J* , 2020 | N/A |
| Control probe for ChIRP (5'-CTCATGGAGTTAGCCCGCTC-3’, 3' biotin-labeled) | Ninomiya et al., *EMBO J* , 2020 | N/A |
| HSATIII-FISH probe (5'-ATTCCATTCCATTCCATTCC-3', 5' Cyanin5-labeled) | Ueno et al, *Mol Cell* , 2026 | N/A |
| PP7-FISH probe ((5’-TTTCTAGGCAATTAGGTACCTTA-3’, 5’-TTTCTAGAGTCGACCTGCAG-3’, 5’-AATGAACCCGGGAATACTGCAG-3’, 5' Cyanin5-labeled) | This paper | N/A |
